# Multi-sample non-negative spatial factorization

**DOI:** 10.1101/2024.07.01.599554

**Authors:** Yi Wang, Kyla Woyshner, Chaichontat Sriworarat, Genevieve Stein-O’Brien, Loyal A Goff, Kasper D. Hansen

## Abstract

Analyzing multi-sample spatial transcriptomics data requires accounting for biological variation. We present multi-sample non-negative spatial factorization (mNSF), an alignment-free framework extending single-sample spatial factorization (NSF) to multi-sample datasets. mNSF incorporates sample-specific spatial correlation modeling and extracts low-dimensional data representations. Through simulations and real data analysis, we demonstrate mNSF’s efficacy in identifying true factors, shared anatomical regions, and region-specific biological functions. mNSF’s performance is comparable to alignment-based methods when alignment is feasible, while enabling analysis in scenarios where spatial alignment is unfeasible. mNSF shows promise as a robust method for analyzing spatially resolved transcriptomics data across multiple samples.

## Background

Spatially resolved transcriptomics (SRT) measures gene expression levels in the context of spatial positions (KH Chen et al., 2015; Ståhl et al., 2016; Rodriques et al., 2019a; Stickels et al., 2021; Y Lee et al., 2021; Zhao et al., 2022; Lubeck, Cai, 2012; Eng et al., 2019; Goltsev et al., 2018; Keren et al., 2019; Thornton et al., 2021), either at the single cell level or as a local aggregate of multiple cells across a spatial location, also termed a spot. The last 30 years of genomics have established that it is essential to consider biological replicates when trying to understand a biological system (Schurch et al., 2016; Mendelevich et al., 2021). Indeed, technology does not remove biological variation (Hansen et al., 2011).

Multisample (population-level) analysis of spatial data is common in functional magnetic resonance imaging (fMRI) brain data, and it is instructive to briefly review the approach in this field (Ombao et al., 2019). In fMRI analysis, the first step is to spatially align the samples to a common coordinate system (known as template-based alignment). The unit of measurements are 3D cubes known as “voxels”. Following alignment, analysis then proceeds separately for each voxel (or sometimes region), typically by using a general linear model across samples. For fMRI data, spatial alignment makes it possible to deploy standard statistical models for each voxel separately, substantially simplifying downstream analysis.

In spatially resolved transcriptomics, a number of methods for spatial alignment has been proposed, including PASTE (Zeira et al., 2022), PASTE2 (Liu et al., 2023), STalign (Clifton et al., 2023) and GPSA (Jones et al., 2023). Some of these methods align to a common coordinate system, others align the samples to each other. However, we posit that there are natural limitations to the potential success of this approach to multi-sample analysis. In fMRI imaging, alignment is helped by the fact that the whole brain is imaged in 3D in each sample. In contrast to fMRI data, the alignment of spatially resolved transcriptomics is complicated by the possibility that different samples may be collected from different anatomical areas and have differences in the shape, size, and rotation of the sections. Indeed, SRT samples can represent completely disjoint areas; in this case, spatial alignment is impossible except to a common coordinate system. But even then, it is unclear how downstream analysis should proceed, when the samples are non-overlapping.

Factor analysis has been a successful approach to unsupervised discovery of patterns in genomics. There are a few existing methods for the factor analysis of SRT data that model the spatial dependency of gene expression data (Townes, Engelhardt, 2023; Velten et al., 2022; Shang, Zhou, 2022b). NSF (Townes, Engelhardt, 2023) and MEFISTO (Velten et al., 2022) are focused on the factorization of data from a single biospecimen, as each factor is modeled using a single Gaussian Process. MEFISTO allows multi-sample analysis, but it requires the samples to be spatially aligned as a preprocessing step. Similarly, NSF can be applied to multiple samples, provided they have been spatially aligned. The spatial alignment process, preceding MEFISTO and NSF analyses, transforms samples to a common coordinate system. This creates an expanded covariance matrix encompassing all samples, with inter-spot correlations based on aligned distances. Both MEFISTO and NSF then utilize this aligned, multi-sample data structure for their analyses. Neither methods are evaluated for this purpose in their respective publications. In contrast, SpatialPCA (Shang, Zhou, 2022b) can be applied to unaligned samples and the publication contains a light evaluation of this task. The authors compares clusters obtained from a joint respectively single-sample analysis to manually annotated cortical layers and conclude that the multi-sample analysis does not outperform single-sample analysis. The SpatialPCA model has a few limitations. Primarily is lacks the ability to have a sample-specific parametrization of spatial dependencies for individual factors, such as the bandwidth in its Gaussian Process components. This restriction potentially hampers its effectiveness when examining samples of diverse sizes or those originating from different platforms. Together, this suggests that across-sample factorization of SRT data still has substantial challenges.

## Results

### Bypassing spatial alignment by parameter modeling

It is important to account for sample-to-sample variation in the analysis of genomics data. We are considering this question in the context of applying matrix factorization methods, such as non-negative matrix factorization (NMF), to spatially resolved transcriptomics data. Gene expression data exhibits a spatial dependence whereby genes measured at two locations which are spatially close show a different dependence from genes measured at two distant spatial locations. Such dependence can be driven by a variety of sources including spatial patterns in the distribution of cell types as well as correlated measurement error. The goal of any analysis of spatially resolved transcriptomics data is to identify systematic changes in gene expression which are associated with spatial location. Broadly, we refer to such dependence as “spatial dependence”.

One approach to analysis of multi-sample spatially resolved transcriptomics data is to start the analysis with spatial alignment of the samples into a common coordinate system. This process essentially defines spatial neighbourhoods and maps these neighbourhoods between samples. But spatial alignment is well-recognized to be a challenging problem, due to the need to account for differences in shape, rotation, and placement of anatomical regions or other features between samples.

Here, we provide a general recipe for extending a one-sample spatial factorization framework to allow multi-sample analysis. Our approach bypasses the need for spatial alignment.

In a spatial factorization framework, we represent spatial expression data on a single sample as a sum of products between gene loadings and spatial factors,

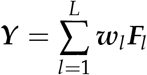

Here ***Y*** is the spatial data matrix, *l* = 1, …, *L* is the number of spatial factors, ***w***_*l*_ are gene loadings and ***F***_*l*_ are the spatial factors. The gene loadings and spatial factors represent systematic changes in gene expression. Accounting for spatial dependence in such a model is done by additional modeling of the spatial factors; an example is the proposed non-negative spatial factorization (NSF) of Townes, Engelhardt (2023) where the spatial factors are modelled using Gaussian processes.

In a multi-sample dataset we have an additional *m* index for the different samples. Our recipe prescribes letting the spatial factors be sample-specific while the gene loadings are shared across samples (Methods), giving rise to the following factorization

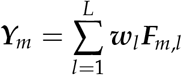

Note the absence of the *m* index on the gene loadings ***w***_*l*_. Sharing the gene loadings across samples is exactly what happens when NMF is applied to data without spatial information such as bulk RNA-seq data (Methods).

With such a parameterization, our recipe enforces each factor to have the same association with genes across samples, while allowing spatial dependence to be modeled separately in each sample.

Fitting such a model will usually require the development of new software (Methods), often by extending existing software.

As a proof of concept, here we have applied this recipe to non-negative spatial factorization (NSF). We allow each sample to have its own spatial dependence structure (or more specifically a sample-specific covariance term in the Gaussian process). We call this extended model mNSF (multisample NSF). We provide a python package implementing our model.

### The performance of mNSF in simulations

We examined the performance of our mNSF model using a simulation study which is a simple extension of the study conducted by Townes, Engelhardt (2023), but adapted to examine issues that are particularly relevant for multi-sample analysis of SRT data. Briefly, we specify true latent spatial factors and generate gene expression data with noise. We only depict the true and estimated spatial features, and not the gene loadings. We use T1-4 to denote the 4 true factors in the simulation, and use M1-4 to denote the 4 mNSF spatial features. There are no particular order to the mNSF output so we manually identify the best true factor which matches a given estimated factor.

First, we examined how mNSF handles the important case where the spatial factors are rotated between samples (T1-T4 in Figure 1a). For each factor, its spatial distributions are the same in the two samples, but rotated 90 degrees. We find the mNSF factors among M1-4 each corresponding to one simulated factor among T1-4, according to their spatial pattern (M1, M2, M3 and M4, corresponding to T2, T1 T4 and T3). We use Moran’s I to measure the spatial dependency for each of the mNSF factors (i.e. M1-4) (Figure 1c), and we find that all four mNSF factors show high spatial dependency. Those validations suggest that mNSF successfully identifies these simulated spatial features.

**Figure 1.**
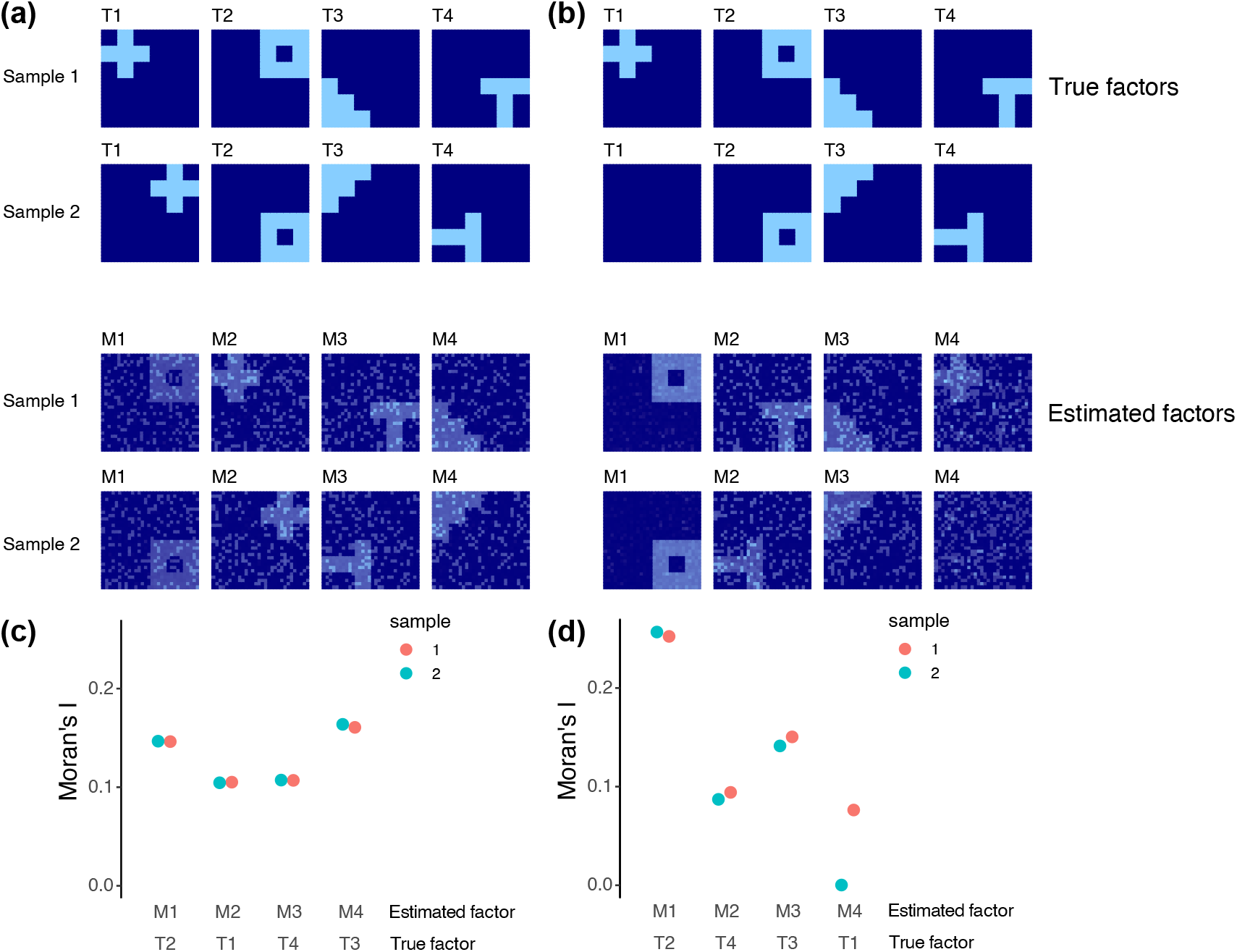
Multi-sample NSF on simulated data. **(a)** We generate 4 true factors for 2 samples, labelled T1-T4. Using these 4 spatial features, we generate noisy expression data for 500 genes following the approach in (Townes, Engelhardt, 2023). Simulated true factors are shown in the top two rows. Factors in sample 2 are the same as sample 1 except they are each rotated 90 degrees anticlockwise. mNSF factors are shown in the bottom two rows. The order of the factors estimated by mNSF is arbitrary and we manually pair each true factor with the estimated factor which best represent it. **(b)** As **(a)** but factor T1 for sample 2 is set to 0. **(c**,**d)** Moran’s I for each of the 4 estimated factors across the 2 samples in the **(c)** first and **(d)** second simulation.

Second, we examined mNSF in the case where one of the factors has a spatial pattern only in one sample, and has constant low value in the other sample (i.e. T1 among the factors T1-T4 in Figure 1b). We find mNSF factors each corresponding to one simulated factor (M1, M2, M3 and M4, corresponds to T2, T4, T3 and T1). For each set of simulations, we use Moran’s I to measure the spatial dependency of each of the mNSF factors among M1-4 (Figure 1d). We find that factor M4, which corresponds to factor T1 (i.e. the factor which is only operational in one of the samples by design), show high spatial dependency in sample 1 and almost zero spatial dependency in sample 2. Those results suggest that mNSF is capable of identifying factors that represent patterns that are operational only in some of the samples.

Third, we examined the performance of mNSF under additional complex conditions, including size differences between samples (Supplementary Figure S1a), pattern distortions between samples (Supplementary Figure S1b), and sample-specific noise levels (Supplementary Figure S1c). These results demonstrate mNSF’s robustness in extracting meaningful patterns despite varying noise, size and shape of the signal across samples.

### Analysis of a mouse sagittal brain dataset

To assess the ability of mNSF to identify spatially-resolved features in an actual, multi-sample, SRT dataset, we next analyzed the adult mouse sagittal brain dataset generated by 10X genomics, generated using the Visium technology (Ståhl et al., 2016). This dataset consists of two sagittal sections from a single mouse brain. Each section was cut in two halves, one anterior and one posterior, for a total of 4 samples. Each sample was assayed using a separate Visium slide. We know that certain anatomical regions are present only in the anterior (e.g. olfactory bulb) or the posterior (e.g. cerebellum), while other anatomical regions will be split across the two halves (e.g. the hippocampus). This provides an opportunity to assess the ability of mNSF to identify sample-specific factors as well as both common factors and factors that vary across the anterior and posterior sections.

Townes, Engelhardt (2023) applies NSF to one of the two anterior samples. They use 20 factors with a split of 10 spatial factors and 10 spatially-unrestricted features. Despite this demarcation, they find that most of the 20 learned patterns have strong spatial components. They establish that most of these factors correspond to known anatomical regions in the anterior mouse brain. Consistent with their parameter choices, we apply mNSF to all 4 samples using 20 spatial features.

To interpret each mNSF factor, we find a list of genes that are highly associated with, and most specific for each factor by analyzing the gene loading matrix using the patternMarkers approach identified in (Fertig et al., 2010). We then use the set of genes associated with each factor to interpret the cell types, biological functions, or anatomical regions represented by each factor.

Some mNSF factors reflect specific anatomical regions, only present in some of the samples. For example, factor M9 can be identified as highlighting the gyri of the cerebellum (Figure 2). This hindbrain-specific factor is close to zero for the two anterior samples and visually similar in intensity and distribution across the two replicate sections for the posterior brain. The top genes identified by patternMarkers include *Pcp2, Calb1, Car8*, and *Itpr1* which are specific markers for Purkinje cells within the cerebellum (X Chen et al., 2022), as well as *Cbln1* and *Cbln3* which are known to exhibit high expression in the cerebellum, and *Zic1* which is a specific marker for cerebellar granular cells (Aruga et al., 1998) (Supplementary Table S1). This result highlights the ability of mNSF to identify factors associated with a signal that is present only in some, but not all, of the samples.

**Figure 2.**
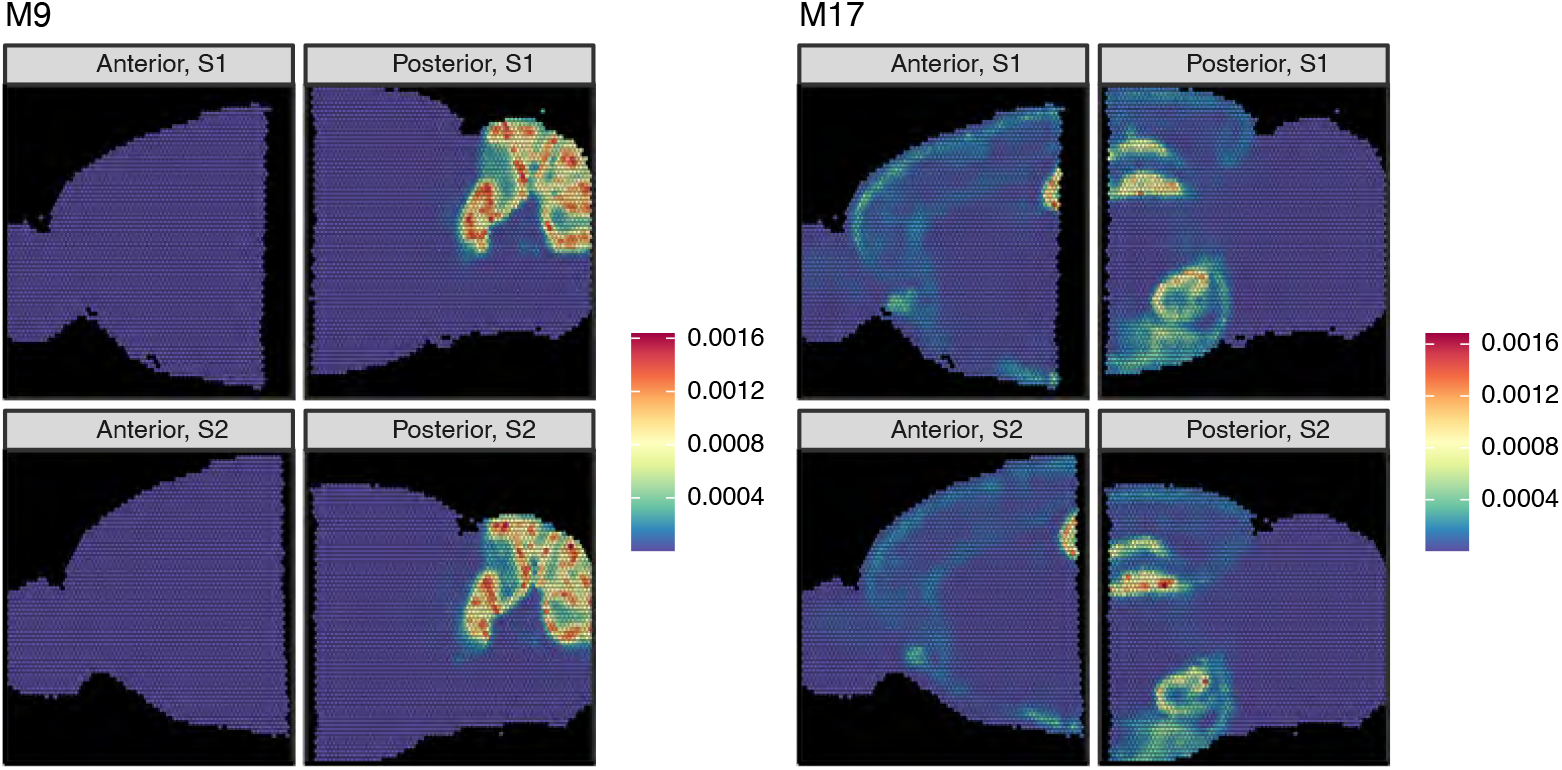
mNSF factors of mouse sagittal data reflect anatomical structures. The dataset is composed of four samples from the same mouse. The sagittal brain was divided into an anterior and a posterior half and two parallel sections were obtained for each half, for a total of 4 samples. We depict the results of applying mNSF using 20 factors to these 4 samples. Factor M9 (left) exhibits high use in the cerebellum and is an example of a factor specific to the posterior brain. Factor M17 (right) exhibits high use in both the hippocampus and hypothalamus regions, where the hippocampus region spans the posterior and anterior areas. This is an example of a factor which varies continuously across disjoint samples. All 20 factors are depicted in Supplementary Figure S2

Some mNSF factors represent anatomical regions that span the posterior and anterior brain (Figure 2). For example, factor M17 predominately marks both the hippocampus and hypothalamus, with moderate signal in select layers of the cortex. Note how the estimated factor varies smoothly across the posterior and anterior brain, specifically across the CA1-3 layers(which appear as a rotated U in these samples). The regions labeled by this factor are considered regions of increased synaptic plasticity. For example, the hippocampus, which functions primarily in learning and memory (Bliss, Collingridge, 1993), requires this plasticity for the formation and consolidation of short-term memories. The hypothalamus plays a crucial role in maintaining homeostasis in the body (Saper, Lowell, 2014), regulating a variety of essential functions such as hunger, thirst, sleep, circadian rhythms, stress responses, and reproductive behaviors. Synaptic plasticity in the hypothalamus is important for adaptation to changes in physiological states and the environment (Dietrich, Horvath, 2013; Serrenho et al., 2019; Bains et al., 2015; Horvath, 2006). Consistent with this categorization, patternMarker genes for M17 are associated with synaptic plasticity, including AMPA receptor regulation (*Cnih2*, Herring et al. (2013)), dendritic spine development (*Ddn, Ncdn*, Yang et al. (2024) and Nicolas et al. (2022)), and synaptogenesis (*Nptxr*, SJ Lee et al. (2017)) (Supplementary Table S1).

Broadly, across the 20 spatial features, our observations about factors M9 and M17 hold for other spatial features. Specifically, we observe (a) consistency between each factor across the two replicate sections (b) there are multiple factors which continuously vary across the anterior and posterior brain (Supplementary Figure S2) (c) a few factors (M10, M19) are specific to either the anterior or posterior brain.

Sensitivity analysis on the dispersion parameter demonstrates that varying the dispersion parameter (ranging from 0.001 to 100) does not significantly affect the factorization results (Supplementary Figures S3, S4). The goodness-of-fit (measured by Poisson deviance between observed counts and predicted mean values) decreases as the number of factors increases, with the decrease rate noticeably slowing after 16 factors (Supplementary Figure S5).

### Analysis of human DLPFC data

Next, to evaluate mNSF across a dataset with replicate samples from different donors, we apply mNSF to a widely used spatially resolved transcriptomics dataset from the Visium platform on human dorsolateral prefrontal cortex (DLPFC) Maynard et al. (2021). This data consists of 4 samples from each of 3 donors. The 4 samples from each donor consist of parallel sections (Figure 3). Each section has a width of 10*µm* and we label the sections as A-D, representing the physical ordering of the sections. The physical separation is as follows: the AB pair is separated by 0 *µ*m, the BC pair is separated by 300 *µ*m and the CD pair is separated by 0 *µ*m. Furthermore, the data are supplied with manual annotations of cortical layers based on expression and H&E staining, with labels of white matter (WM), cortical layers 1 to 6 and NA (which are excluded from our analysis). Not all layers are present in all samples; specifically, the 4 samples from individual 2 do not have layers 1 and 2 present.

**Figure 3.**
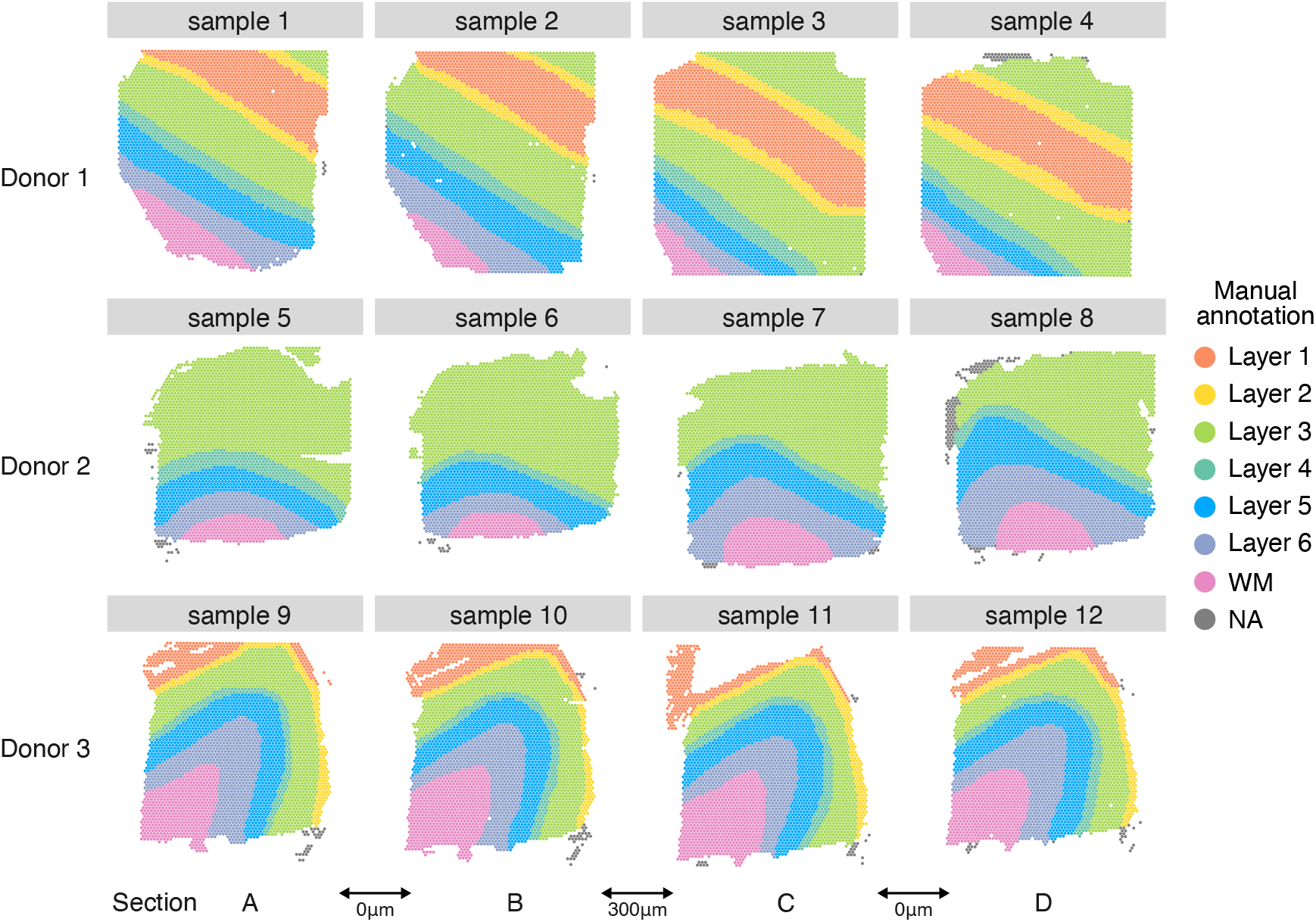
Layer annotation of DLPFC data. Manually annotated layer for each spatial location in each sample from each donor in DLPFC dataset. Each donor contributes 4 different sections, arbitrarily labelled A-D. For samples from donor one and three, each sample has seven layers: layer 1-6 and white matter. The samples from donor 2 do not cover layers 1 and 2. While the manual annotation includes the NA (not available classification), these spatial locations are excluded from our analyses.

We apply mNSF to all 12 samples using 10 spatial features. The model does not encode the design of the experiment with 4 sections from 3 donors. We expect that different samples from the same donor are more similar than different samples from different donors; this is true for the manual annotations of the samples (Figure 3). We do not necessarily expect that different cortical layers form distinct “clusters” in expression space. For example, it is understood that some genes are expressed in a gradient across the cortical layers (O’Leary et al., 2007; Lau et al., 2021; Lodato, Arlotta, 2015). Nevertheless, we expect some relationship between cortical layers and mNSF spatial features. To compare the mNSF factors with the discrete manual annotation, we use the following approach: we group each spatial location in each sample according to its manual annotation and display each factor value across the layers (Figure 4). The M6 factor displayed in Figure 4 is particularly high in cortical layer 2, followed by blending into cortical layer 3. The lowest layer is cortical layer 1 and the factor is almost absent in white matter.

**Figure 4.**
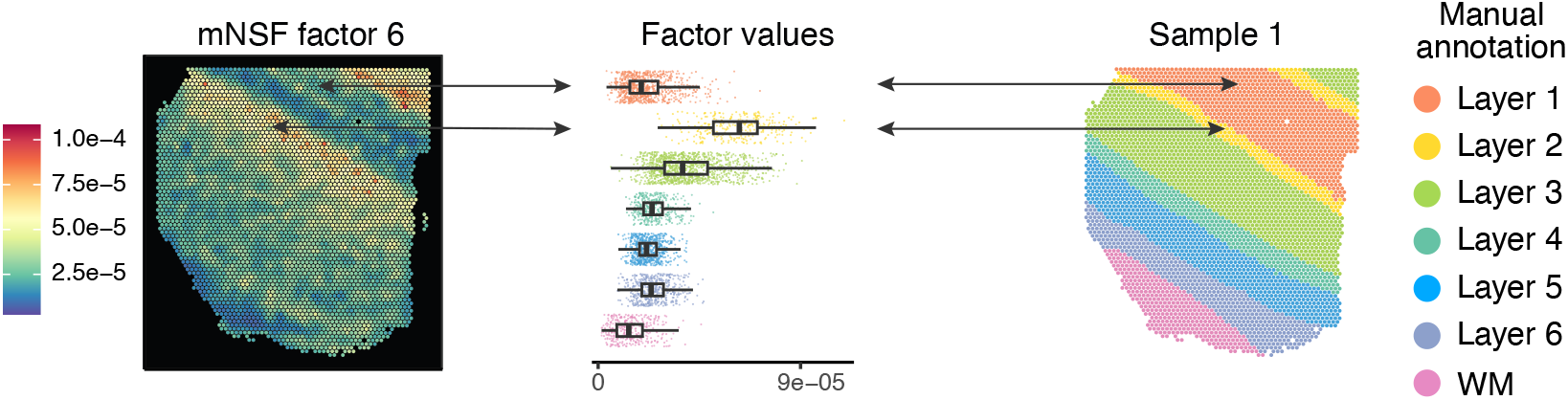
Cross-layer trend of the M6 factor in sample 1. The left plot depict the spatial distribution of factor M6 on the spatial locations from sample 1 (donor 1, section A). The middle plot depicts the distribution of this factor, stratified by which layer each spatial location belongs to (according to the manual layer annotation). The right plot depicts the manual annotation for this sample. The color scale in the left plot is the factor values whereas the color in the middle and left plot is used to depict the manual annotation.

Figure 5 depicts 3 mNSF spatial features. Due to size restrictions, we display 4 samples, 1 sample from each donor as well as an additional parallel section from each donor. This display depicts both between-donor variability and between-sections-within-a-donor variability. For completeness, we depict all 12 samples and 10 factors in Supplementary Figures S6-S15.

**Figure 5.**
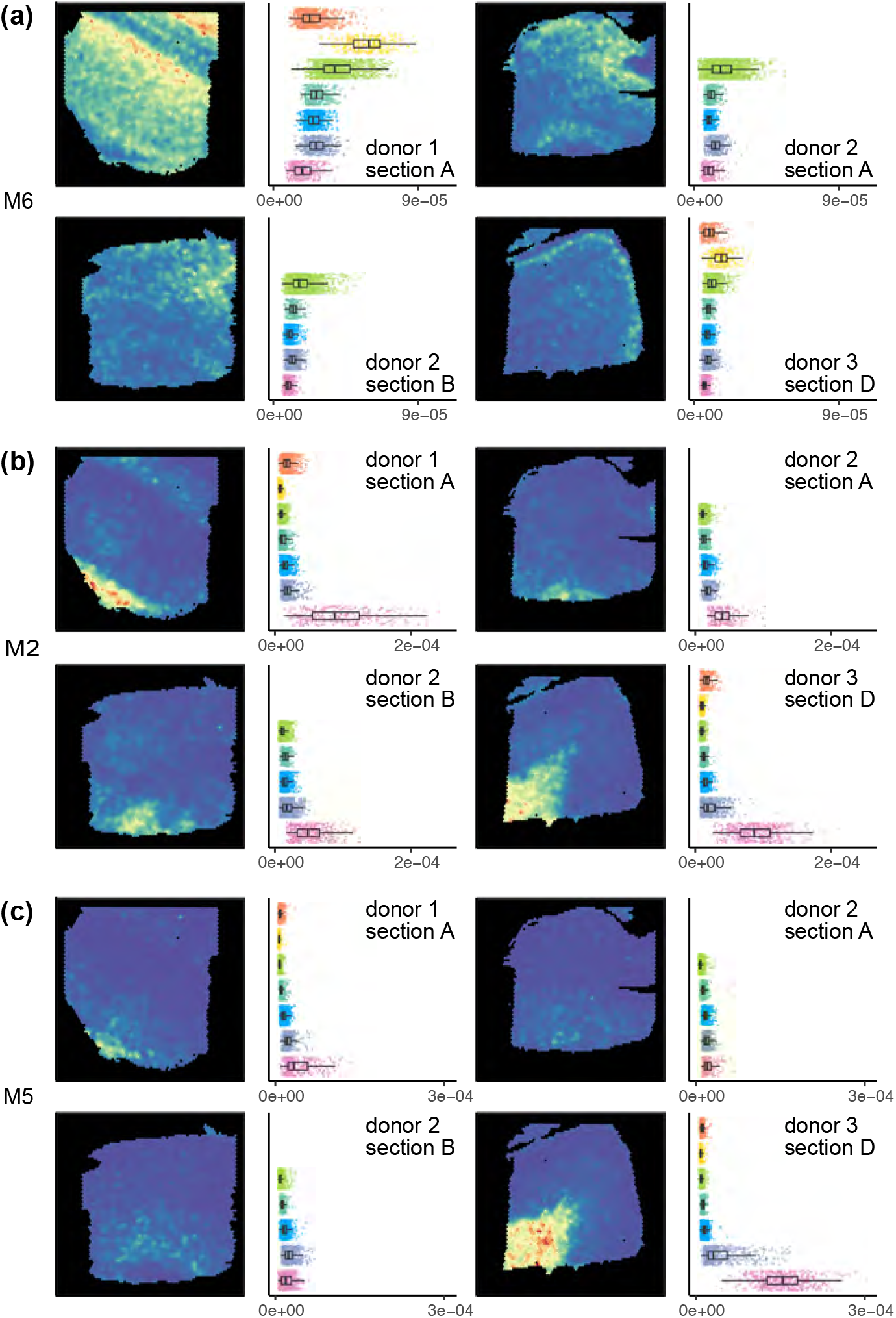
mNSF factors represent layers of the prefrontal cortex. mNSF **(a)** factor M5, **(b)** factor M2 and **(c)** factor M6 in sample 1 (donor 1, section A), sample 5 (donor 2, section A), sample 7 (donor 2, section B) and sample 12 (donor 3, section D).

Factor M6 has high values in the spatial locations manually annotated as layer 2, intermediate values in spatial locations manually annotated as layer 3 and close to zero values for spatial locations manually annotated as white matter (Figure 5a). PatternMarker genes for this factor include HPCAL1 (Supplementary Table S2), which is a marker for layer 2 excitatory neurons (Wei et al., 2022). This factor is consistent with the manual annotation across the 3 donors, and across parallel sections within each donor.

Factor M2 is consistently high for spatial locations annotated as white matter (Figure 5b) and low otherwise. PatternMarker genes for this factor include known oligodendrocyte and myelin-associated genes, such as MOG, MOBP, MBP, and BCAS1 (Cahoy et al., 2008; Plant et al., 2014) (Supplementary Table S2), which are expected to be specific to white matter. This factor is consistent with the manual annotation across the 3 donors, and across parallel sections within each donor.

Factor M5 has high values in the spatial locations manually annotated as white matter in the samples from donor 1 and 3, but consistently low values across all the spatial locations in the samples from donor 2 (Figure 5c). It therefore suggests an inconsistency, potentially in tissue processing or orientation, across the samples from different donors. PatternMarker genes for this factor include genes that mark oligodendrocytes and myelination (PLP1, TF, CNP, ENPP2) (Cahoy et al., 2008; Plant et al., 2014), and genes potentially associated with neurovasculature, blood, and vascular endothelial cells (HBA2, HBB, CLDND1) (Günzel, Yu, 2013) (Supplementary Table S2). We believe differences in this factor across donors may reflect variation in how tissue blocks were dissected where sections from donor 2 are cut at a more horizontal plane that does not contain layers 1 and 2. This was confirmed to be a possible interpretation by the original manual annotator (K. Maynard, personal communication). This factor represents a pattern only present in some, but not all samples, once again showcasing the ability of mNSF to identify such patterns.

Sensitivity analysis of the dispersion parameter demonstrated that factorization results remained stable across a range of values from 0.001 to 10. However, significant changes were observed for dispersion values between 10 and 100 (Supplementary Figure S16). Additionally, the association between mNSF factors and manually annotated layers exhibited strong correlation for dispersion parameters up to 10 but weakened considerably for values larger than 100 (Supplementary Figure S17). This analysis suggests that the factors identified by the model are robust across a wide range of dispersion parameter values.

To address the choice of the number of factors, we evaluated mNSF performance on DLPFC data using 4-20 factors, guided by three metrics: (a) factor-layer association (b) adjusted Rand index (ARI) for domain identification (c) goodness-of-fit (Poisson deviance). We observed increasing cluster stability as more factors are included.(Supplementary Figure S18). We chose to use 10 factors for our analysis of the DLPFC dataset, as this provides a good balance between model complexity and performance.

We conclude that mNSF shows encouraging performance on this dataset. It produces factors which make biological sense and are consistent across parallel sections within the same tissue block. Many of the factors are also consistent across the donors. However, the scale of the factors sometime vary (see white matter for factor M2, Figure 5). This might reflect variability in the manual annotation, biological variability between samples or unwanted (technical) variation which might be possible to remove with additional normalization.

### Comparison to spatial alignment

The DLPFC dataset is an excellent candidate for spatial alignment. However, considering the manual annotations (Figure 3) it is clear that aligning different sections from the same donor is much easier than aligning different sections from different donors. For example, layers 1 and 2 are absent in donor 2 and this complicates spatial alignment. Zeira et al. (2022) describes PASTE, a method for spatial alignment, and apply PASTE to the DLPFC dataset to perform direct alignment of these samples. However, for exactly the challenges described above, Zeira et al. (2022) only attempt to align samples from the same donor to each other, doing both pairwise and 4-sample alignment. Using these data allows us to compare mNSF to spatial alignment on a dataset which is particularly well suited for spatial alignment.

Specifically, we compare the performance of mNSF and PASTE-NSF on identical sample pairs. This approach, which we term pasteNSF, involves using PASTE to align samples followed by NSF on the aligned samples. Following Zeira et al. (2022), we focus on pairwise alignment, considering only adjacent section pairs in our evaluation. Specifically, both methods are applied to the same set of two adjacent section pairs (either AB, BC, or CD) for each donor. For instance, when analyzing the BC pair from Donor 1, mNSF utilizes factors specific to the BC-Donor1 case, while PASTE-NSF employs its alignment followed by NSF, also using factors specific to the BC-Donor1 case. By conducting the comparison in this manner, we ensure a direct and fair assessment of both methods’ effectiveness on the same data.

We expect, and this is confirmed by the authors, that PASTE performs best with adjacent tissue sections separated by 0*µm* (ie. comparisons AB or CD). In this analysis we use 10 spatial spatial features. Following factorization, we use a multinomial model to predict the 5 or 7 manually annotated layers (depending on the donor) as a function of the 10 estimated spatial features, and we use the model fit to assess performance. The model fit measures the association between the 10 inferred factors and the manual annotation.

This approach shows that mNSF has comparable performance to pasteNSF (Figure 6a-b). PASTE supplies the user with a mapping score which represents how well the spatial alignment is performed (higher is better). The pair with the lowest mapping score (donor 1, BC) is the pair where mNSF outperforms pasteNSF the most (Figure 6c-d).

**Figure 6.**
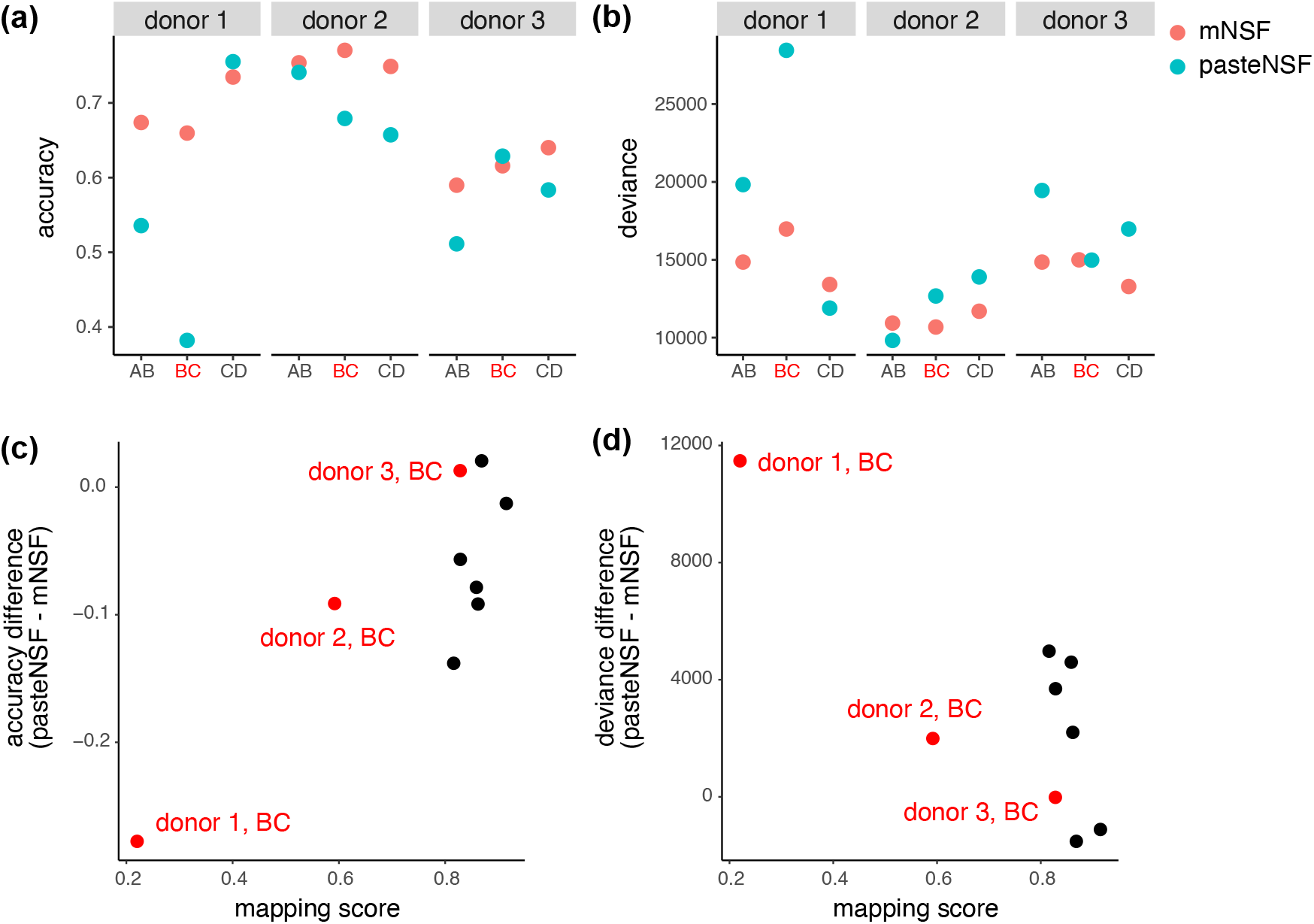
A comparison of mNSF to spatial alignment using PASTE. The performance of mNSF compared with applying NSF on the aligned coordinates by PASTE. Each consecutive sections (within donor) (i.e. sections AB, sections BC or section CD) are used to fit either mNSF, or a combination of PASTE alignment and NSF, using three spatial features. For the factor sets inferred from each pair of samples in each method, we run a multinomial regression of the layer as a categorical variable, against the three spatial features, and calculated the **(a)** the accuracy of layer prediction using a multinomial model (Methods, higher is better). **(b)** The multinomial model deviance (lower is better, Methods). **(c)** Difference in the layer prediction accuracy between mNSF and the combination of PASTE and NSF, compared to the mapping score of PASTE. **(d)** Difference in the deviance of the multinomial regression between mNSF and the combination of PASTE and NSF, compared to the mapping score of PASTE. The BC sections, which are separated by 300 *µ*m, are highlighted in red.

In summary, mNSF has at least comparable performance to PASTE followed by NSF when spatial alignment is easy, but extends factorization to data where spatial alignment is hard (between donors in the DLPFC dataset) or impossible (between anterior/posterior sections in the mouse sagittal dataset).

### Comparison with SpatialPCA

We performed a comparative evaluation of domain identification capabilities between mNSF and SpatialPCA using the DLPFC dataset, with the manually annotated layers providing ground truth. Both methods used 10 factors for consistency. Following the SpatialPCA tutorial, we used walktrap clustering to cluster the continuous factors into discrete sets (Pons, Latapy, 2005; Shang, Zhou, 2022a). Using these discrete sets we measured the performance using the Adjusted Rand Index (ARI) (Figure S19), with ARI values predominantly within the 0.30-0.45 range. Our results show comparable proficiency in identifying spatial domains within this dataset.

Next, we compared spatialPCA and mNSF at domain identification using data across two different technologies, Visium and Slide-seq. We used mouse brain samples containing the cerebellum, from two different datasets. The datasets included a Visium-based mouse sagittal brain dataset focusing on the posterior half (posterior S1, previously described) and Slide-seq data of mouse cerebellum obtained from the Broad Single Cell Portal (ID SCP354) (Rodriques et al., 2019b), with processed data as reported in the SpatialPCA manuscript Shang, Zhou (2022b). This brain region has a very distinctive shape and location which allow us to roughly assess whether the domain is consistent across the samples, even in the absence of a known ground truth. Both methods used 10 factors. We then employed k-NN clustering for domain identification, which requires pre-specification of the number of domains. Using 5 domains, mNSF roughly identifies the cerebellum in the two samples, whereas spatialPCA fails at this task (Figure 7). However, the performance of the mNSF is sensitive to the pre-specified number of domains. Using more domains than 5, the visual agreement between Visium and SlideSeq is not as striking as depicted in Figure 7. In contrast, we were unable to obtain decent results using SpatialPCA despite varying the number of domains. This comparison suggests that mNSF’s have better performance in identifying biologically relevant spatial domains across different technological platforms.

**Figure 7.**
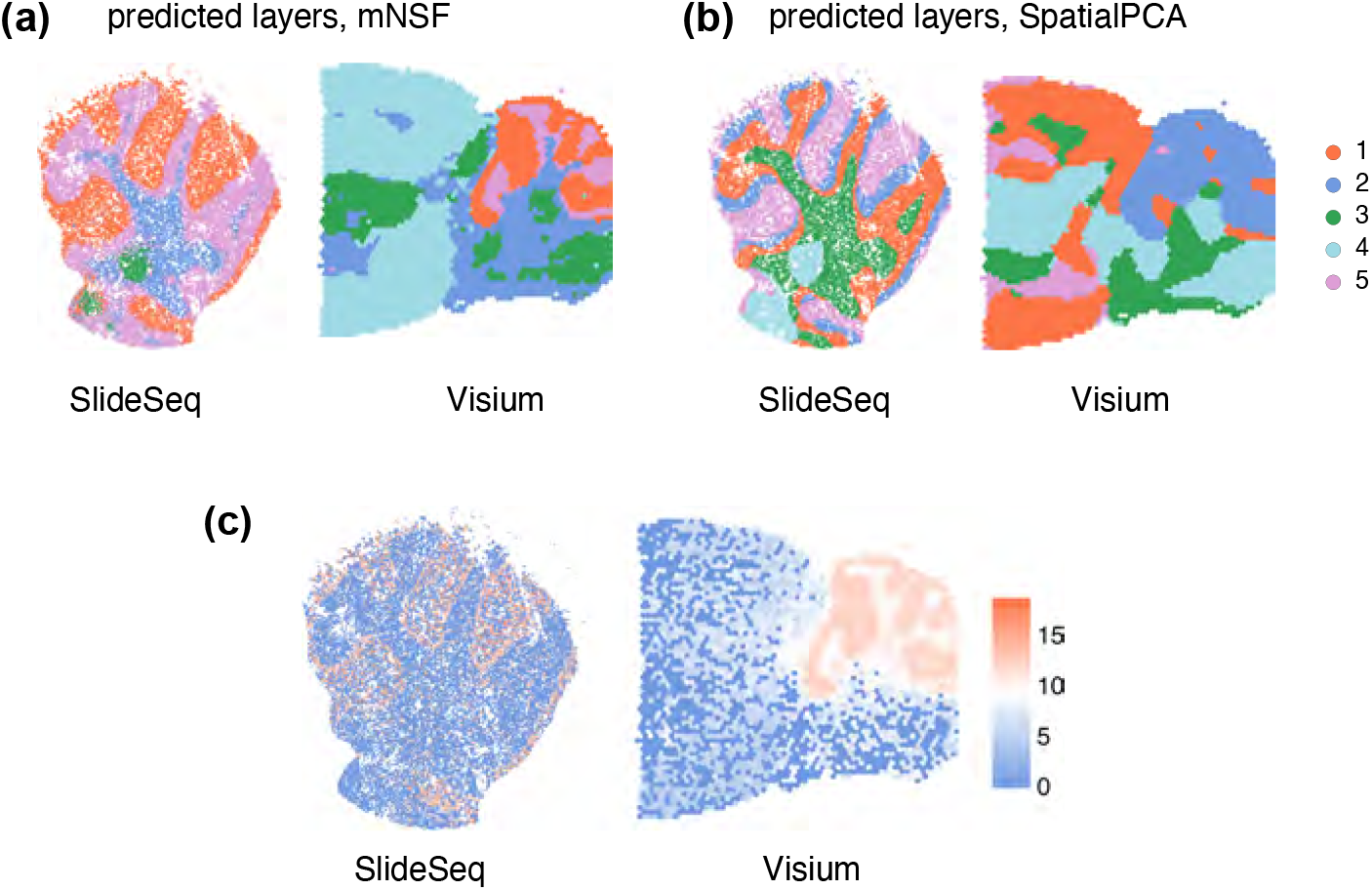
Comparison of domain identification performance between mNSF and SpatialPCA in cross-platform data analysis of mouse cerebellum. Predicted layers by **(a)** mNSF and **(b)** SpatialPCA for the joint analysis of Slide-seq and Visium data. **(c)** Expression level of Pcp2, marker gene of cerebellum in each dataset.

### Memory and time consumption

To investigate memory and time consumption, we ran mNSF multiple times on datasets of various sizes (Supplementary Figure 8) using a GPU. The memory usage has a linear relationship with the number of factors, genes, samples and a quadratic relationship with the number of spots within each sample. Run time has a linear relationship with the number of factors, genes and samples and a quadratic relationship with the number of spots within each sample. The number of genes and the number of factors only has a small impact on run time.

Resource consumption will grow linearly with larger sample size and quadratically with higher resolution. In addition, mNSF is built on tensorflow and is therefore designed to be used on GPUs which have lower memory limits than CPUs. For these reasons, we implemented two strategies to reduce (primarily) memory consumption. The first is the use of inducing points, which was already part of the NSF method. Inducing points is a standard approach used with Gaussian processes: instead of using all data points, they use a smaller set of points for prediction. The second we call chunking and it consists of making multiple passes over the data, each time only using a random set of points. In each pass over random sets of points, we keep the gene weights unchanged (Methods). Chunking and inducing points can be combined. We examined how computational resources scale with the percentage of induced points and number of chunks used in the analysis (Supplementary Figure 8). Memory usage increases linearly with the percentage of induced points, while runtime shows a moderate increase. When increasing the number of chunks, memory usage shows a clear decrease, demonstrating that chunking can be an effective strategy for reducing memory requirements. However, this comes with a trade-off of increased runtime, which rises with the number of chunks.

To understand how model performance scales with these computational parameters, we evaluated mNSF using different percentages of induced points and numbers of data chunks. Using the DLPFC dataset as a benchmark, we assessed both layer prediction accuracy and deviance (Figure S20). The results show that accuracy remains relatively stable down to 25% induced points before declining, while chunking the data into more pieces gradually reduces accuracy. This suggests a trade-off between computational efficiency and model performance. We caution that this trade-off is likely to be dataset dependent.

To examine factor stability under different computational settings, we calculated correlations between factors obtained using different numbers of data chunks (Figure S21). While factors remain generally consistent with 2-4 chunks, using 10 chunks leads to decreased correlations, particularly for some factors. This indicates that excessive chunking may compromise the model’s ability to capture certain spatial patterns.

We performed a similar analysis varying the percentage of induced points used (Figure S22). The correlation matrices demonstrate that most factors remain highly stable even when reducing induced points to 25%, with correlation values typically above 0.8. However, further reduction to 5-10% induced points results in decreased factor stability for some patterns. This analysis helps guide parameter selection by showing where computational savings can be achieved while maintaining factor stability.

The memory and time consumption for the analysis used in this paper is in Supplementary Table S3.

## Discussion

In this study, we describe a general approach to extending a matrix factorization method to multisample datasets. Using this approach, we extended non-negative spatial matrix factorization (NSF) by Townes, Engelhardt (2023) to spatial transcriptomics datasets with multiple samples. Our model allows for a sample-specific spatial dependence structure. Our method bypasses the need to align factors between samples into a consensus coordinate system, which is a challenging problem. Both real and simulated data analysis support that the method yields usable results when applied to data from multiple sources, even if it is impossible to perform spatial alignment. Classic matrix factorization methods are widely used in expression analysis and it is well recognized that it is hard to identify the biological or technical process(es) associated with each factor or pattern. Our method retains this limitation.

There are multiple possible downstream applications of our method, including spatial domain detection. These applications are left for future work. Our evaluation is focused on comparing factors to known anatomical regions, and we have not considered the impact on downstream analyses. Nevertheless, we believe that our method can serve as a foundation or input to downstream analysis of multi-sample data.

Batch effects could cause differences in spatial patterns between samples. Such differences would appear as factors which are variable across samples. Our current method cannot distinguish batch effects from biological variation. It will be an important question for future research to appropriately model and correct batch effects in spatially resolved transcriptomics data.

In applications, researchers are sometimes aware of existing patterns or factors across samples, either at sample level or at the level of spatial features. Accounting for such known biology will require the application of a semi-supervised matrix factorization method. Such methods have been suggested for other analysis domains (Haddock et al., 2022). We believe it will be important to develop such models for spatially resolved transcriptomics data and it is likely that our framework will allow for the extension of such models to multiple samples.

## Conclusions

Here, we provide an alignment-free framework for generalizing a one-sample spatial factorization model to multi-sample data. In simulations, our method is capable of identifying spatial structures which are rotated between samples, as well as structures which only appear in some, but not all, samples. Using real data, we show our method is capable of identifying matched functional regions in multi-sample spatial transcriptomics data.

## Methods

### A general approach to multi-sample spatial factorization

Spatially resolved transcriptomic (SRT) data for a single sample can be represented as a matrix ***Y*** = (*y*_*g,i*_) of expression measures indexed by genes *g* with associated spatial (physical) location ***x*** = (*x*_*i*_). We use the index *i* to index the spatial locations.

Consider a standard non-negative matrix factorization model applied to SRT data on a single sample:

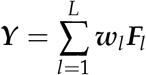

In this model, we decompose the expression values into a term ***w***_*l*_ representing genes and a term ***F***_*l*_ representing the spatial locations. The model does not impose any kind of spatial structure on ***F***_*l*_. This model is widely used for non-spatially resolved bulk and single-cell transcriptomic data. Adapting this model to spatial data is usually done by additional requirements on the ***F***_*l*_ terms to account for expected spatial dependence. Such extensions are considered below, but for the sake of clarity, we first consider a matrix factorization model without spatial dependency.

Our suggested approach to extend this model across *M* samples is to use the following (simplified) model

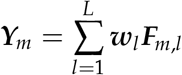

where the gene loadings are shared across samples, but the (spatial) factors *F*_*m,l*_ are sample-specific. The model is easy to fit using standard software for non-negative matrix factorization models, by concatenating the involved matrices:

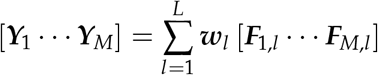

where [·] is concatenation. This is possible because of the simple model formulation where we do not impose spatial structure on the spatial features.

As argued by Townes, Engelhardt (2023), this model could be improved by (a) incorporating the digital (discrete) nature of the expression data and (b) modelling the spatial dependence between spots.

As a first step towards a better model for SRT data, Townes, Engelhardt (2023) describes probabilistic NMF (PNMF) which models the discrete nature of digital expression data using a Negative Binomial distribution but does not address the spatial dependence. We propose a multi-sample version of PNMF (mPNMF) specified as

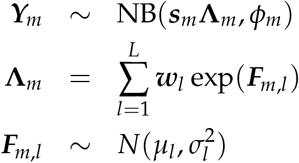

Here, ***s***_*m*_ is a vector of known sample-specific size factors (one for each spot), and 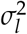 are factorparameters which are not sample-specific. Any software that fits single-sample PNMF can fit multi-sample PNMF by concatenating the data matrices.

To model the spatial dependence of SRT data, Townes, Engelhardt (2023) develops non-negative spatial factorization (NSF) by using a Gaussian process to model the spatial factors ***F***_*l*_. We propose a multi-sample version of this model (mNSF), which is stated as follows (Figure 9):

**Figure 8.**
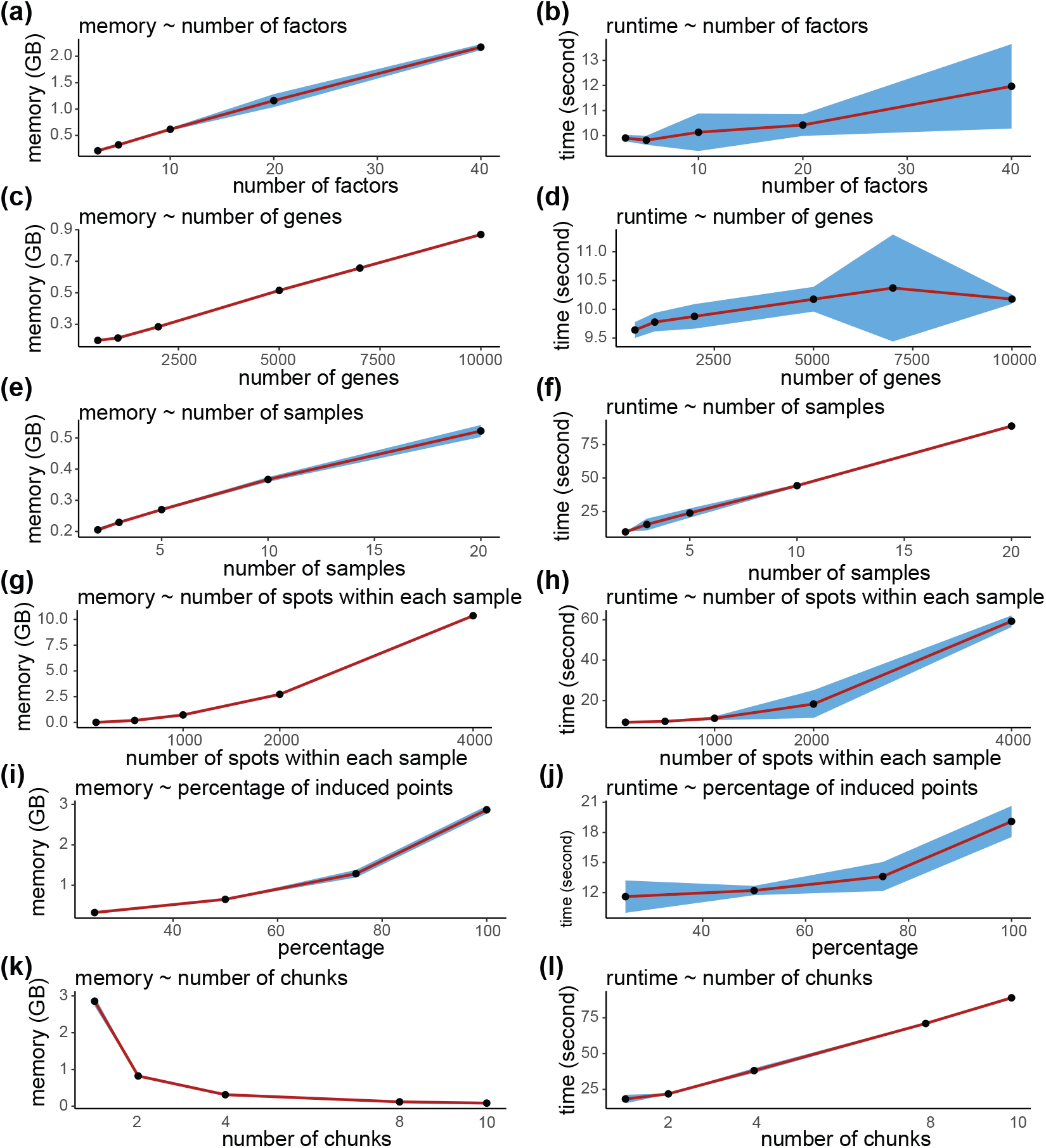
Memory and time profile of mNSF. **(a,b)** Memory usage and runtime vs. number of factors. **(c,d)** Memory usage and runtime vs. number of genes. (e,f) Memory usage and runtime vs. number of samples. **(g,h)** Memory usage and runtime vs. number of spots within each sample. Each experiment is repeated three times. Points represent the average memory usage (or runtime) across three runs. The blue shaded area indicates the 95% confidence interval based on 10 training sets.

**Figure 9.**
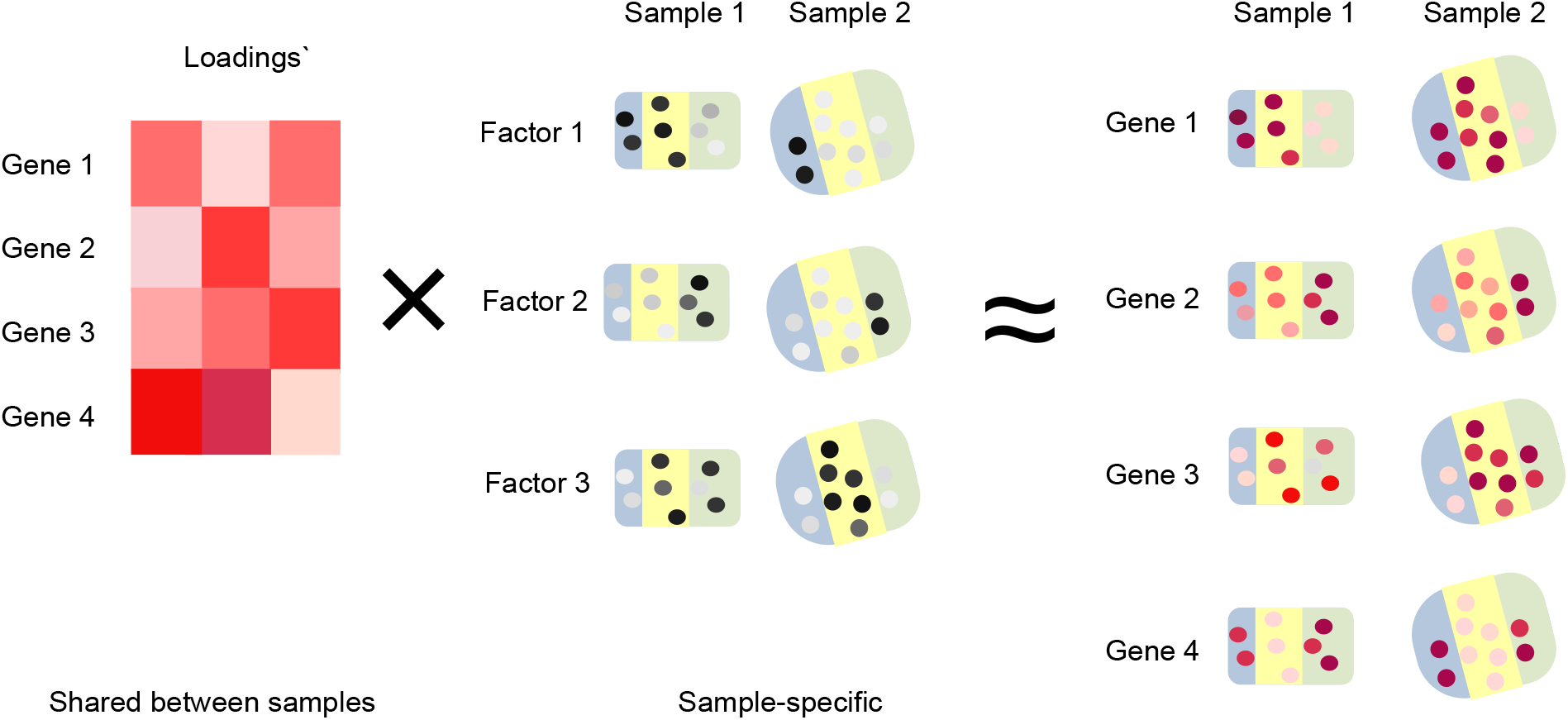
Multi-sample NSF model. Multi-sample NSF model for a dataset with two samples and three spatial features. As input, we use expression data from the two samples where rows are genes and columns are spatial spots. Each factor is modeled by a Gaussian process which represents the spatial dependency between spots; this process has sample-specific parameters. The gene loadings are shared between the two samples.

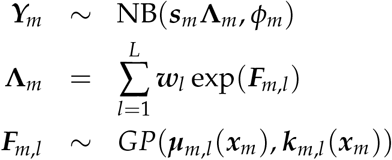

where ***µ***_*m,l*_ is a sample- and factor-specific mean function and ***k***_*m,l*_ is a sample- and factor-specific covariance kernel, both depending on the sample-specific vector of spatial locations ***x***_*m*_. Unlike the multi-sample version of PNMF, fitting mNSF requires sample-specific parameterization to handle the sample-specific spatial features.

We have implemented mNSF by extending the code provided by Townes, Engelhardt (2023). Our extension includes the interpolated version of NSF, where the model is fit using a subset of spatial locations which are then interpolated to encompass the entire data matrix.

### Data processing

#### Mouse sagittal section data

The spot-level gene expression counts data, as well as the 2-dimensional coordinates denoting the position of each spot, are downloaded from 10X website: https://cf.10xgenomics.com/samples/spatial-exp/1.1.0/V1_Mouse_Brain_Sagittal_Anterior_Section_1, https://cf.10xgenomics.com/samples/spatial-exp/1.1.0/V1_Mouse_Brain_Sagittal_Anterior_Section_2, https://cf.10xgenomics.com/samples/spatial-exp/1.1.0/V1_Mouse_Brain_Sagittal_Posterior_Section_1, and https://cf.10xgenomics.com/samples/spatial-exp/1.1.0/V1_Mouse_Brain_Sagittal_Posterior_Section_2.

Top 500 genes are selected based on the maximal Poisson deviance of each gene across all four samples, calculated by a built-in function in NSF package (see code on GitHub for details).

#### DLPFC data

DLPFC dataset are downloaded from the SpatialExperiment (Righelli et al., 2022) package. Top 500 genes are selected based on the maximal Poisson deviance of each gene across all twelve samples (see code on GitHub for details).

#### PASTE alignment

For each sample pair used in this study, spatial locations on the pairwise aligned coordinate system are downloaded from PASTE GitHub website: https://github.com/raphael-group/paste_reproducibility

The 500 genes selected in the 12-sample mNSF analysis for DLPFC data are used for this analysis.

For each sample pair used in this study, one-sample NSF is applied on the aligned data, i.e. the concatenated gene expression matrix of the two samples as well as the coordinate of each spatial location in each sample on the aligned coordinate system. mNSF is then used on the unaligned data, i.e. the gene expression matrix of each sample as well as the coordinate of the spatial locations in each sample in the original coordinate system.

### Models without induced points

#### One-sample NSF

As reference, we describe the one-sample NSF model proposed in (Townes, Engelhardt, 2023). Briefly, the model assumes that the log value of each non-negative factor follows a Gaussian process across the spatial locations in the sample, with the intercept equal to a linear combination of the coordinates. The gene expression level in each spatial location follows a Poisson distribution with the mean equal to the product of a loading matrix and the factor matrix, multiplied by the size factor (i.e. library size) of the spot.

Assume there are *n* spatial locations in total at locations **X** measuring *G* genes. We will use *L* to denote the number of non-negative spatial spatial features; this is a user-supplied parameter.

Let *Y*_*gi*_ denote the observed count value for gene *g*^*th*^ and spatial location *i*. It is assumed to follow a Negative Binomial distribution

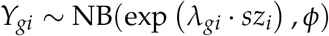

Here *sz*_*i*_ is a known size factor for spatial location *i, ϕ* is the dispersion parameter and

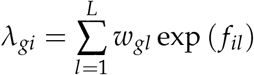

Here *w*_*gl*_ denotes the loading of gene *g* for the *l*^*th*^ factor, and *f*_*il*_ denotes the value of the *l*^*th*^ factor at spatial location *i*.

The value of *l*^*th*^ factor on the observed spatial locations ***X*** follows a Gaussian process distribution with a linear mean and a Matern kernel for covariance. In particular, this means that ***F***_*l*_(***X***) – the specification of the factor on the grid ***X*** follows a normal distribution

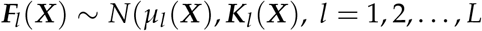

with

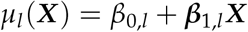

and

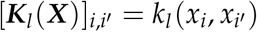

where the kernel function for the *l*^*th*^ factor is of Matern class,

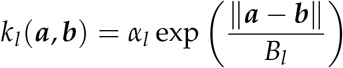

Here

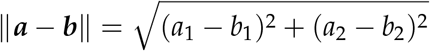

denotes the distance between two spatial locations with coordinates ***a*** and ***b***, both of which are vectors of length 2.

In summary, ***K***_*l*_ is an *N × N* matrix denoting the correlation of ***F***_*l*_, *α*_*l*_ is the length scale parameter and *B*_*l*_ is the amplitude parameter for the kernel of Gaussian process.

#### Multi-sample NSF

If no interpolation is used, multi-sample NSF assumes that the log value of each non-negative factor follows a Gaussian Process across the spatial locations in each sample, with the intercept equals a linear combination of the coordinates. And the gene expression level in each spatial location follows a Poisson distribution with the mean equals a weighted sum of the factors multiplied by the size factor (i.e. library size) of the spot, where the weights are shared across different samples and the other parameters are all sample-specific.

Assume there are *N*_*m*_ spatial locations in total at locations **X**_**m**_, *G* genes used, and *L* non-negative spatial spatial features.

The observed count value for *g*^*th*^ gene at *i*^*th*^ spatial location in the *m*^*th*^ sample, denotes as *Y*_*mgi*_, follows a Negative Binomial distribution

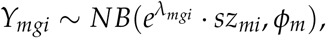

where *sz*_*mi*_ is the scale factor of spatial location *i, ϕ*_*m*_ is the dispersion parameter, and

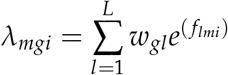

Here *w*_*il*_ denotes the loading of gene *j* for the *l*^*th*^ factor, and *f*_*lmi*_ denotes the value of the *l*^*th*^ factor at spatial location *i* in the *m*^*th*^ sample.

The value of *l*^*th*^ factor on the observed spatial locations X conditional on ***U***_*lk*_ follows a GP distribution

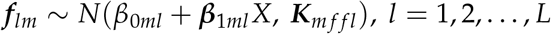

Here ***K***_*m f f l*_ is an *N*_*m*_ × *N*_*m*_ matrix denoting the correlation of ***f***_*lm*_, with

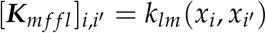

where the kernel function for the *l*^*th*^ factor in sample *m* is

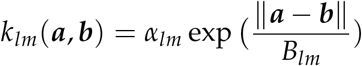

where *α*_*lm*_ is the length scale parameter and *B*_*lm*_ is the amplitude parameter for the kernel of Gaussian Process for the *l*th factor in the *k*^*th*^ sample.

### Models with inducing points

#### One-sample NSF

If a set of interpolated points is used, one-sample NSF assumes that the log value of each non-negative factor follows a Gaussian Process across both the observed and interpolated spots, with the mean equals a linear combination of the coordinates. A set of parameters are created for the interpolated points, and the posterior distribution of the observed point conditional on the interpolated points is derived. The overall likelihood of both the observed and interpolated points is calculated through the likelihood of the interpolated points and the posterior likelihood of the observed points.

Assume there are *N* spatial locations in total at locations **X**, *J* genes used, *n* spatial locations interpolated at locations ***Z***, and *L* non-negative spatial spatial features.

The observed count value for *g*^*th*^ gene at *i*^*th*^ spot, denotes as *Y*_*gi*_, follows a Negative Binomial distribution

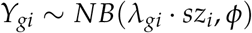

where *sz*_*i*_ is the scale factor of spatial location *i, ϕ* is the dispersion parameter, and

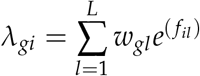

Here *w*_*il*_ denotes the loading of gene *j* for the *l*^*th*^ factor, and *f*_*il*_ denotes the value of the *l*^*th*^ factor at spatial location *i*.

The distribution of ***U***_*l*_ (i.e. the value of the *l*th factor on the induced points) and ***F***_*l*_ follows a Gaussian Process distribution,

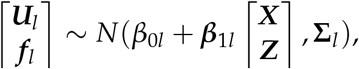

where

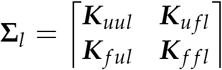

Here ***K***_*uul*_ is an *n* × *n* matrix denoting the correlation of ***U***_*l*_, ***K*** _*f f l*_ is an *N* × *N* matrix denoting the correlation of ***F***_*l*_, and ***K***_*u f l*_ is an *n* × *N* matrix denoting the correlation between ***U***_*l*_ and ***f***_*l*_.

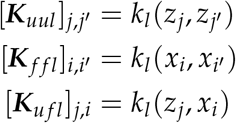

where the kernel function for the *l*^*th*^ factor is

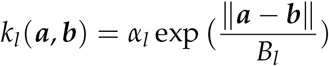

where *α*_*l*_ is the length scale parameter and *B*_*l*_ is the amplitude parameter for the kernel of Gaussian Process for the *l*th factor.

Decomposing the joint distribution of ***U***_*l*_ and ***F***_***l***_ into *P*(***U***_*l*_) and *P*(***F***_*l*_ | ***U***_*l*_), we have

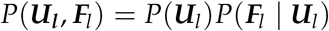

where *P*(***U***_*l*_) could be derived by

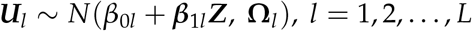

and *P*(***F***_*l*_ | ***U***_*l*_) could be derived by

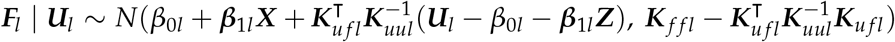

#### Multi-sample NSF

In multi-sample NSF, for each factor, the loading of the same gene is shared across the samples, while all the other parameters are sample-specific. The observed data from different samples are assumed to be independent.

Assume there are K samples, with the *m*^*th*^ sample containing *N*_*m*_ spatial locations at ***X***_*m*_, *n*_*m*_ interpolated points at ***Z***_*m*_. The same set of *G* genes are used in all the samples. Assume there are L non-negative spatial factors for each sample, with the loadings of those *G* genes for each factor shared by samples.

For sample *m*, the observed count value for *g*^*th*^ gene at *i*^*th*^ spot, denotes as *Y*_*mgi*_, follows a Negative Binomial distribution

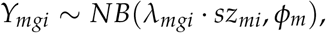

where *sz*_*mi*_ is the scale factor of spatial location *i* in sample *m, ϕ*_*m*_ is the dispersion parameter of sample *m*, and

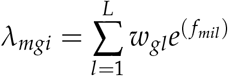

Here *w*_*gil*_ denotes the loading of gene *j* for the *l*^*th*^ factor for sample *m*, and *f*_*mil*_ denotes the value of the *l*^*th*^ factor at spatial location *i* in sample *m*.

The value of the *l*^*th*^ factor on the interpolated locations of sample *k* are assumed to follow a GP distribution

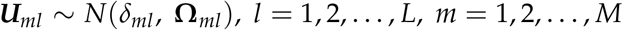

The distribution of ***U***_*ml*_ and ***F***_*ml*_ follows a Gaussian Process distribution,

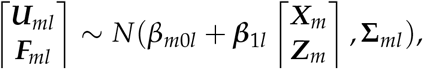

where

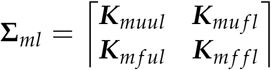

Here ***K***_*muul*_ is an *n*_*m×*_ *n*_*m*_ matrix denoting the correlation of ***U***_*ml*_, ***K***_*m f f l*_ is an *n*_*m*_ *×n*_*m*_ matrix denoting the correlation of ***f***_*ml*_, and ***K***_*mu f l*_ is an *n× N* matrix denoting the correlation between ***U***_*lm*_ and ***f***_*lm*_.

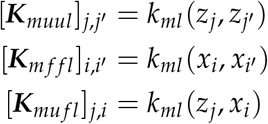

where the kernel function for the *l*^*th*^ factor in the *m*^*th*^ sample is

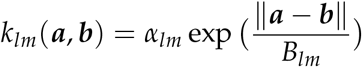

where *α*_*lm*_ is the length scale parameter for sample *m* and *B*_*lm*_ is the amplitude parameter for the kernel of Gaussian Process for sample *m* for the *l*th factor.

Decomposing the joint distribution of ***U***_*lm*_ and ***F***_*lm*_ into *P*(***U***_*lm*_) and *P*(***F***_*lm*_ | ***U***_*lm*_), we have

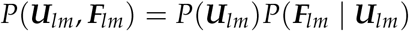

where *P*(***U***_*lm*_) could be derived by

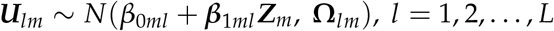

and *P*(***F***_*lm*_ | ***U***_*lm*_) could be derived by

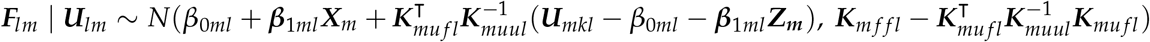

#### Model fitting

Firstly, let’s assume a one-sample data with the distribution in the same form of one-sample NSF, as described in the first subsection under **Method** section, and discuss it’s model fitting approach.

For one-sample spatial data, in NSF paper, it has been shown that by maximizing the following function (called ELBO function), we will get the MLE estimates of all the parameters involved in the model (Townes, Engelhardt, 2023):

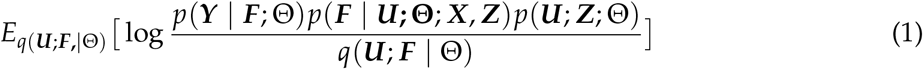

where Θ denotes the parameter space, ***F***[, *l*] is defined by letting ***F***[, *l*] = ***f***_*l*_, and *q*(***U, F*** | Θ) is the product of the posterior likelihood of ***F*** conditional on ***U***, denoted as *q*(***F*** | ***U, X, Z***, Θ), and the approximated likelihood of ***U***, denoted as *q*(***U*** | ***Z***).

Next, we will discuss the model fitting approach for multi-sample data, where the distribution of the data is in the same form of the mNSF model.

The statement that “maximizing the ELBO function will give us the MLE estimates of all parameters involved in the model” is hold in general regardless of the form of distribution settings, such statement also holds for a data that is concatenated by data from multiple samples, where each data has the same form of distribution but with different values of parameters, i.e.

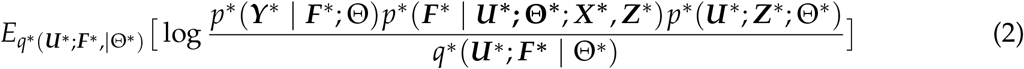

with

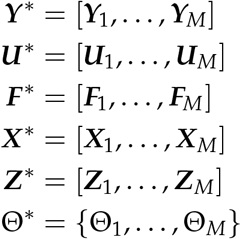

and where ***Y***_*m*_ is the observed data at all spatial locations in sample *m*, ***U***_*m*_ denotes the latent factors at induced points in sample *m*, ***F***_*m*_ is the factor at all spatial locations in sample *m*, ***X***_*m*_ is the spatial locations in sample *m*, and ***Z***_*m*_ is the induced points in sample *m*.

As discussed in the last paragraph, the statement “maximizing the ELBO function will give us the MLE estimates of all parameters involved in the model” holds true for function (2), so in the next step, we will discuss the approach to maximize the function (2) above.

One way to maximize function (2) is using “Adam algorithm (Kingma and Ba, 2014) with gradients computed by automatic differentiation in Tensorflow” (Townes, Engelhardt, 2023), which calculate the gradient of a target function with respect to a set of parameters and update the parameters by adding *s*· *g* to each of the parameter where *s* denote the ‘step size’ (a constant scalar that has the same value for fitting different parameters) in the gradient approach and *g* denotes the gradient of a parameter. To satisfy the non-negativity constraint of *W* parameter, we can set any negative values in *W* to zero after the parameters’ update in each iteration.

In the setting that the distributions of data from different samples are independent, we can re-write function (2) as

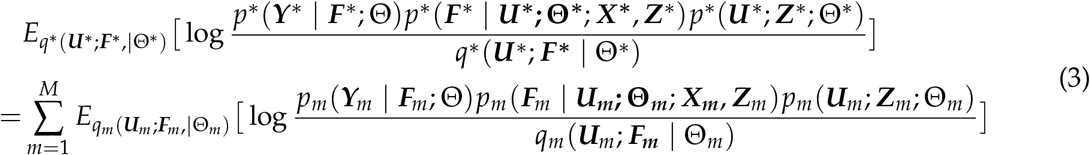

The equation (3) above suggests that, in terms of the gradient calculation and the parameters update within each iteration of applying Adam gradient approach in the multi-sample model fitting, it equals to

Step 1: calculate the gradient of the parameters involved in each sample, only using the data of the corresponding sample;

Step 2: for the parameters that are sample-specific, update those parameters in the same way of fitting one-sample NSF model; for the parameters that are shared by samples (here for mNSF model, it is the loadings parameter ***W***), the gradient of this parameter for function (3) equals the sum of the gradients of the parameter across *m* samples.

Note that as long as the ‘step size’ parameter are the same for the individual sample’s model fitting, the sample-specific parameter fitting in “Step 2” equals:

Step 2*: for the sample-specific parameters, update those parameters separately in the same way of fitting one-sample NSF model, (here in mNSF model, we will get *M* sets of updated *W*s, written as *W*_*m,new*_), then average those updated parameters to get the updated parameter with respect to the full model (here for mNSF model, the updated *W* parameter can be calculated by 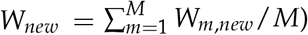

Based on all the discussions above in this subsection, we can draw the conclusion that the following two model fitting process will give us the same parameter estimates:

Process 1: maximize the ELBO function of the full mNSF model, using Adam gradient approach with step size of *s* and updating the parameters with 100 iterations, where at the end of each iteration, set the non-negative values in the averaged ***W*** to zeros.

Process 2: repeat the following parameter updating step for up to 1000 iterations, until converge: for each sample: firstly do the parameter updates in the same way as one iteration in ‘Process 1’ excluding the step of setting the negative values in ***W*** to zero; then for parameter ***W***, get the average of its updated value across the *M* samples, set the non-negative values in the averaged ***W*** to zeros, and use this non-negative ***W*** as the updated ***W***

In mNSF, we use “Process 2” to fit the model, which will, assuming the approximated likelihoods used in NSF model fitting are close enough to the non-approximated likelihoods, give us a estimate of parameters that is close to the MLE estimates of the model.

#### Data chunking implementation

Mini-batch processing, where data is divided into smaller subsets for parallel processing, has become ubiquitous in deep learning due to its efficiency in handling large datasets while maintaining model convergence. Drawing inspiration from this approach, mNSF implements data chunking for analyzing large spatial transcriptomics datasets. This approach divides the spatial locations into multiple chunks and processes them in parallel, providing another layer of computational efficiency on top of the inducing points method.

In our implementation, the spatial locations in each sample are first randomly partitioned into several chunks. Each chunk contains a subset of the original spatial locations. The number of chunks can be specified by the user based on their computational resources and dataset characteristics. This chunking is performed independently for each sample.

The model is then fit to each chunk separately while maintaining the shared gene loadings across all chunks and samples. Each chunk only needs to store and process the corresponding subset of the data. The results from different chunks are then combined during the parameter update step of the model fitting process. Importantly, the gene loadings remain shared across all chunks and samples, ensuring that the biological interpretation of the factors remains consistent across the entire dataset.

The distributed points approach can be used in combination with inducing points, providing two complementary strategies for handling large datasets.

#### Parameter selection for induced points and data chunks

The inducing points approach, implemented in the NSF by Townes, Engelhardt (2023), provides an efficient approximation method for handling spatial correlations. Our implementation of mNSF extends this approach while adding data chunking as a complementary strategy for computational efficiency.

While our implementation of mNSF provides both induced points and data chunking for computational efficiency, selecting appropriate parameters for these approximations requires careful consideration. The optimal parameters depend on both computational constraints and data characteristics. The key trade-off is between the number of induced points (U) and number of chunks (K) per sample, as both parameters affect memory usage and computational efficiency, with induced points primarily impacting model accuracy and chunks affecting parallel processing capability.

In our cross-platform analysis of mouse brain data (Visium sagittal brain and Slide-seq cerebellum), we used different parameter settings for each technology due to their distinct characteristics. For the Visium data (N=3,355 spots), we selected 35% of spatial locations as induced points and processed the data as a single chunk. For the larger Slide-seq dataset (N=25,415 spots), we maintained the same proportion of induced points but divided the data into 10 chunks. This parameter selection was guided by multiple considerations: the cerebellum’s distinctive layered structure requires sufficient induced points to capture its complexity, while the large number of Slide-seq spots benefits from parallel processing through chunking.

For the analyses of both the DLPFC dataset (12 samples, ∼3,500 spots per sample) and the mouse sagittal brain dataset (4 samples, 2,500-4,000 spots per sample), we selected 35% of spatial locations as induced points and processed each dataset as a single chunk. This parameter choice balances computational efficiency with model accuracy. The relatively moderate size of these datasets (less than 5,000 spots per sample) and their well-defined anatomical structures (cortical layers in DLPFC, distinct brain regions in sagittal sections) made this configuration suitable. The 35% induced points provided sufficient coverage to capture the spatial patterns while maintaining reasonable computational demands. Single-chunk processing was appropriate given the manageable sample sizes and the importance of preserving spatial relationships within each sample.

These parameters should be adjusted based on specific dataset characteristics. Datasets with fine spatial structures (e.g., cortical layers) may require more induced points while potentially using more chunks. In contrast, datasets with large homogeneous regions may achieve good results with fewer induced points. When samples vary greatly in size or complexity, consider sample-specific parameters. Users should validate parameter choices by comparing results with different settings on a subset of data, monitoring convergence behavior, and assessing whether anatomically meaningful patterns are preserved.

## Availability of data and materials

The Visium data for DLPFC is available for download through SpatialExperiment package. The Visium data for mouse sagittal section is available through 10X portal (https://www.10xgenomics.com). Code for generating the aligned spatial coordinates using PASTE is available through GitHub (https://github.com/raphael-group/paste_reproducibility). All code to analyze the data and generate figures is available at https://github.com/hansenlab/mNSF_paper and Zenodo (DOI:10.5281/zenodo.13881001). Our mNSF package is available at https://github.com/hansenlab/mNSF. Example codes for using mNSF are publicly available at https://github.com/hansenlab/mNSF/blob/main/tutorial/mnsf-tutorial-dlpfc.md and https://github.com/hansenlab/mNSF/blob/main/tutorial/mnsf-tutorial-mouse.md.

This work is licensed under GNU Lesser General Public License v3.0.

## Declarations

## Funding

Research reported in this publication was supported by the National Institute of General Medical Sciences of the National Institutes of Health under award number R35GM149323 (YW, KDH), the National Institute on Aging of the National Institutes of Health under award numbers R01AG066768 (KW, CS, LAG), R01AG072305 (LAG), the National Institute of Neurological Disorders and Stroke of the National Institutes of Health under award number R00NS122085 (CS, GSO), the National Cancer Institute of the National Institutes of Health under award number U01CA284090 (GSO), the Raynor Foundation (GSO) and the Chan Zuckerberg Initiative DAF, an advised fund of the Silicon Valley Community Foundation under award CZF2019-002443 (YW, KDH).

## Conflicts of Interest

None.

## Author contributions

YW developed the method and performed analyses, supervised by KDH. KW, CS, GSO and LAG provided feedback on the method, software, evaluation and interpretation of the method. YW and KDH wrote the manuscript, with feedback and help from all other authors.

## SUPPLEMENTARY MATERIALS

### Multi-sample non-negative spatial factorization

Yi Wang, Kyla Woyshner, Chaichontat Sriworarat, Genevieve Stein-O’Brien, Loyal A Goff, Kasper D. Hansen

## 1 Supplemental Tables

**Supplementary Table S1.**
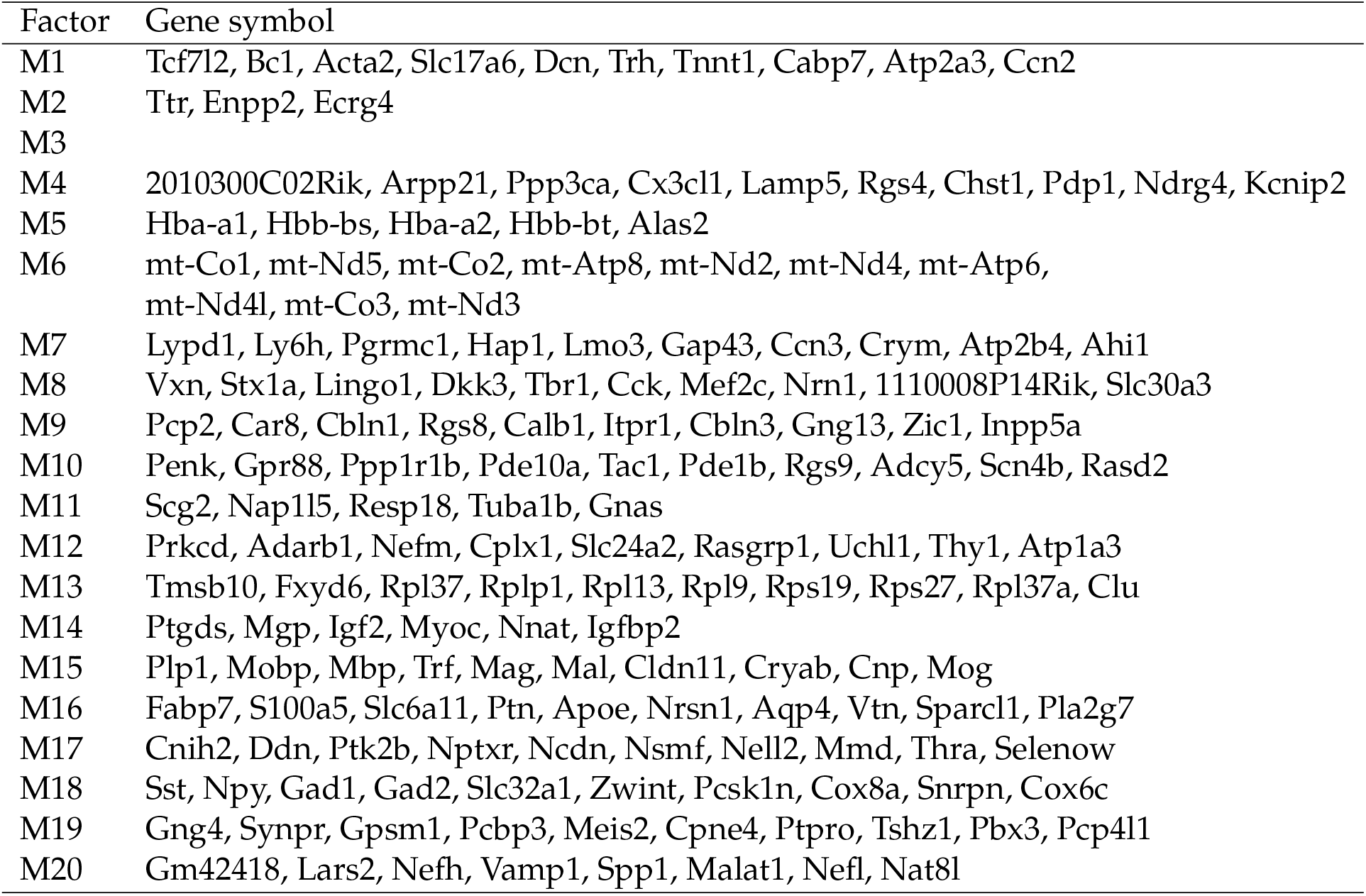
Genes mostly associated with each factor in mouse sagittal section data |.

**Supplementary Table S2.**
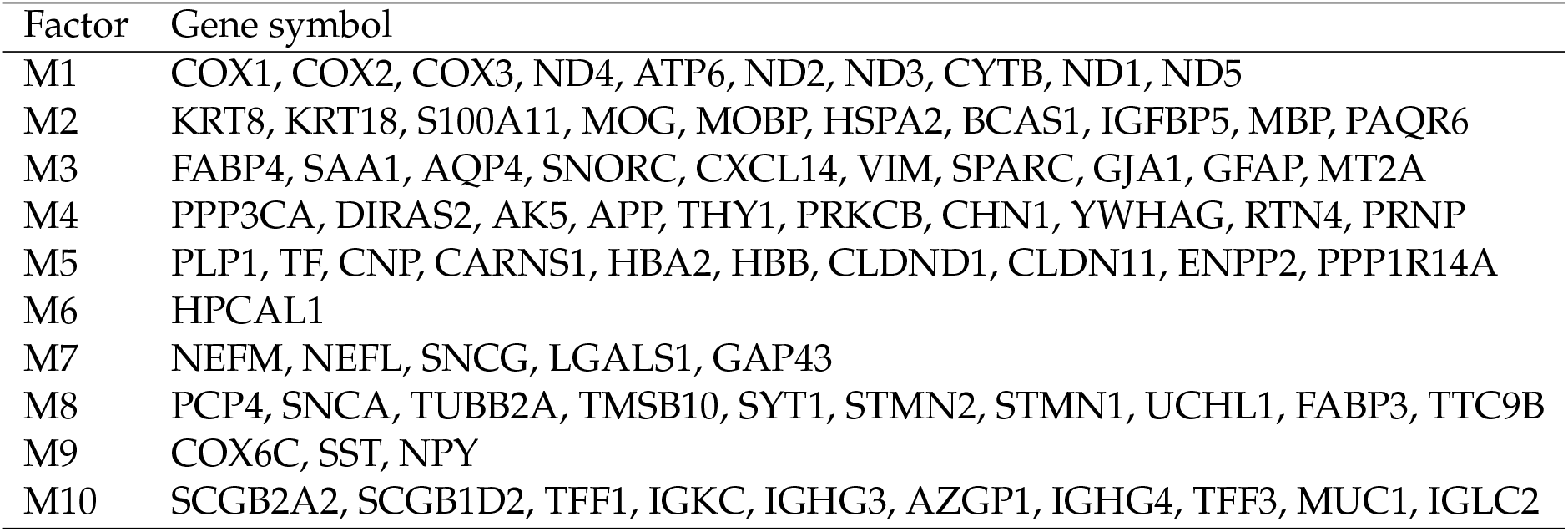
Genes associated with each factor in the DLPFC data |.

| Factor | Gene symbol |
| --- | --- |
| M1 | COX1, COX2, COX3, ND4, ATP6, ND2, ND3, CYTB, ND1, ND5 |
| M2 | KRT8, KRT18, S100A11, MOG, MOBP, HSPA2, BCAS1, IGFBP5, MBP, PAQR6 |
| M3 | FABP4, SAA1, AQP4, SNORC, CXCL14, VIM, SPARC, GJA1, GFAP, MT2A |
| M4 | PPP3CA, DIRAS2, AK5, APP, THY1, PRKCB, CHN1, YWHAG, RTN4, PRNP |
| M5 | PLP1, TF, CNP, CARNS1, HBA2, HBB, CLDND1, CLDN11, ENPP2, PPP1R14A |
| M6 | HPCAL1 |
| M7 | NEFM, NEFL, SNCG, LGALS1, GAP43 |
| M8 | PCP4, SNCA, TUBB2A, TMSB10, SYT1, STMN2, STMN1, UCHL1, FABP3, TTC9B |
| M9 | COX6C, SST, NPY |
| M10 | SCGB2A2, SCGB1D2, TFF1, IGKC, IGHG3, AZGP1, IGHG4, TFF3, MUC1, IGLC2 |

**Supplementary Table S3.**
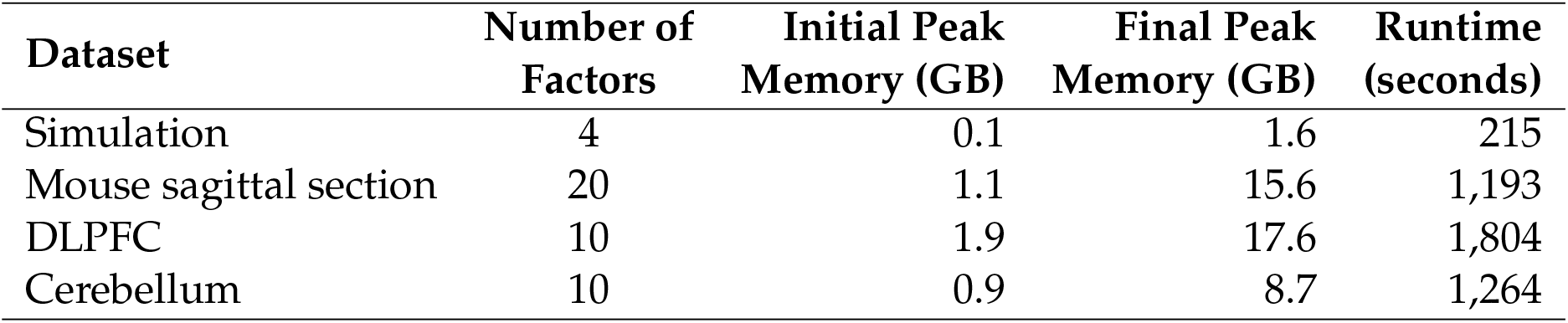
Memory consumption and runtime analysis for different spatial transcriptomics datasets. The table shows the memory requirements at different stages of the analysis pipeline and the total computational time required for each dataset.

## 2 Supplemental Figures

**Supplementary Figure S1.**
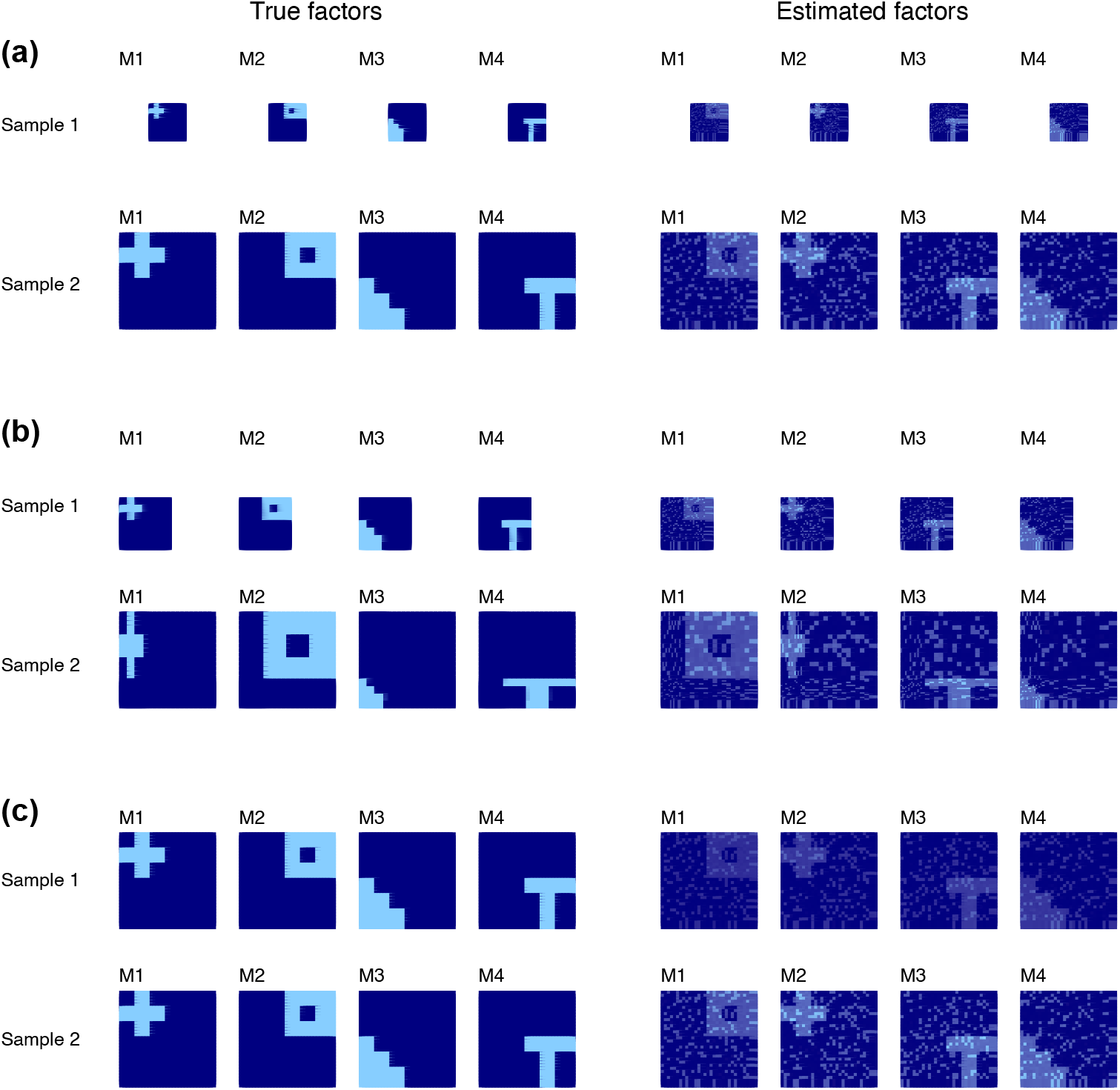
Multi-sample NSF performance on simulated data under additional scenarios. Like Figure 1 but under additional scenarios. True factors are in the first column, estimated factors in the second. **(a)** Size differences: samples with varying sizes of spatial patterns. **(b)** Distortion: the true factor is distorted between samples. (c) Sample-specific noise level.

**Supplementary Figure S2.**
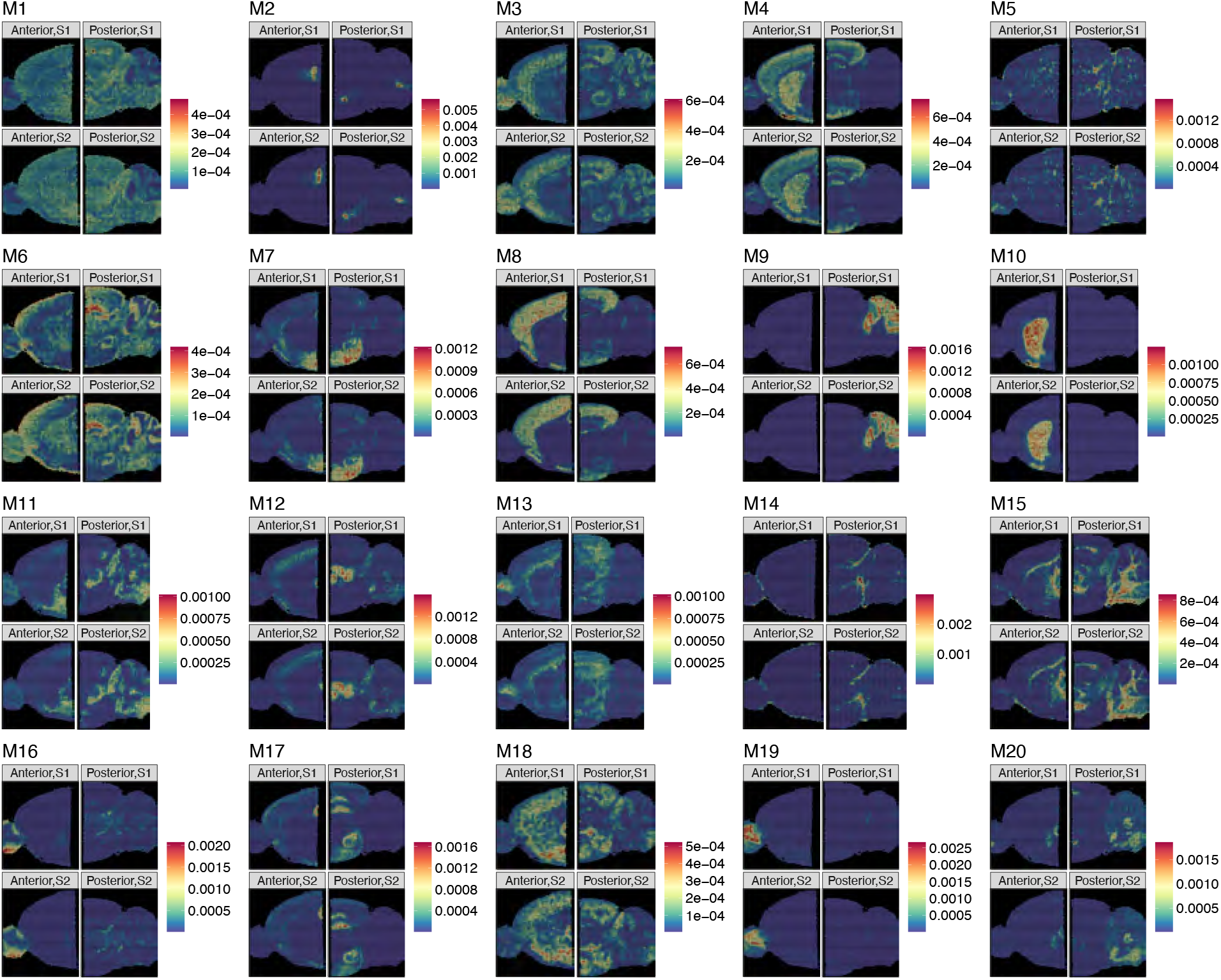
mNSF factors of mouse sagittal data show associations with the anatomical structure. |The dataset is composed of four samples – two pairs of replicates, each for the anterior and the posterior region. Four-sample NSF is applied in this data, with twelve factors used. Each pair of replicates is in the same column in each subplot. Comparing the spatial pattern of each factor to a reference diagram of the mouse brain, it is easy to establish that factor 16 and 19 are enriched in olfactory bulb, and factor 9 is enriched in cerebellum.

**Supplementary Figure S3.**
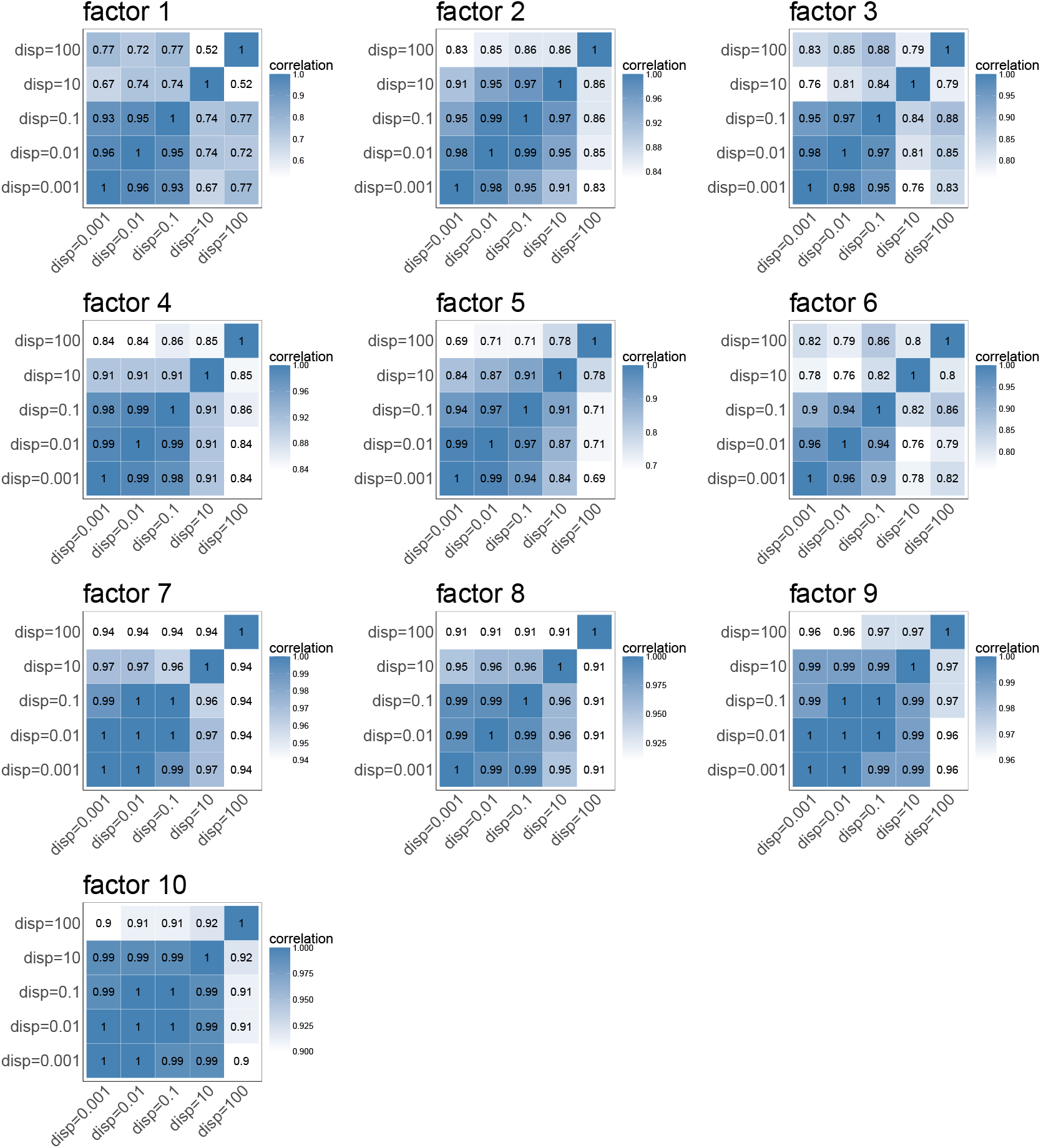
Sensitivity analysis of dispersion parameter on mouse sagittal brain data. Correlation matrices displaying the similarity between spatial factors obtained using different dispersion parameter values (ranging from 0.001 to 100) for factors 1-10 in the mouse sagittal brain dataset. Each heatmap represents one factor, with correlation values indicated by color intensity. The consistently high correlation values (generally > 0.8) across most dispersion parameter values demonstrate that mNSF produces stable results across a broad range of dispersion values (0.001, 0.01, 0.1, 10, and 100). Some factors show slightly lower correlations at dispersion=100.

**Supplementary Figure S4.**
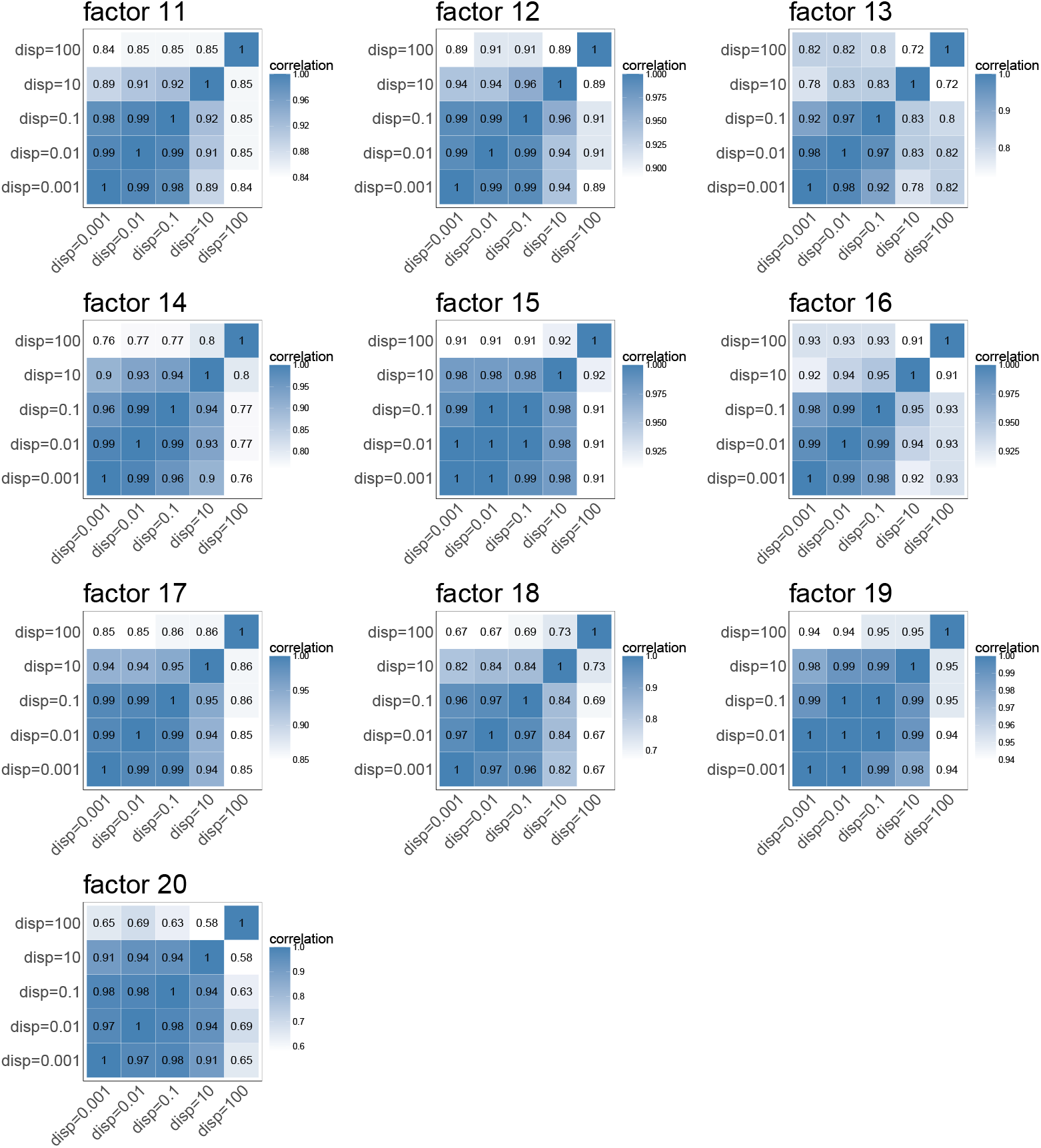
Sensitivity analysis of dispersion parameter on mouse sagittal brain data. Like Supplementary Figure S3 but for factors 11-20.

**Supplementary Figure S5.**
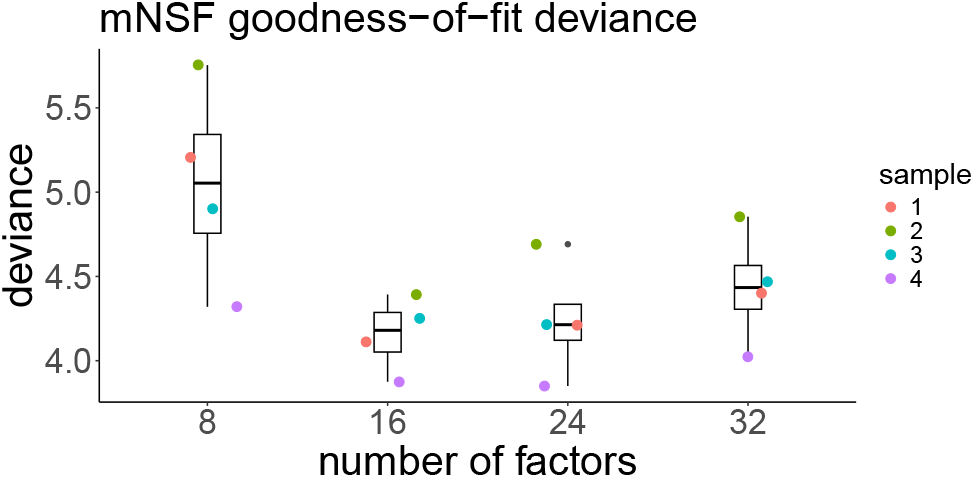
Evaluation of mNSF performance across different numbers of factors for mouse sagittal section data. Goodness-of-fit deviance for mouse sagittal section data.

**Supplementary Figure S6.**
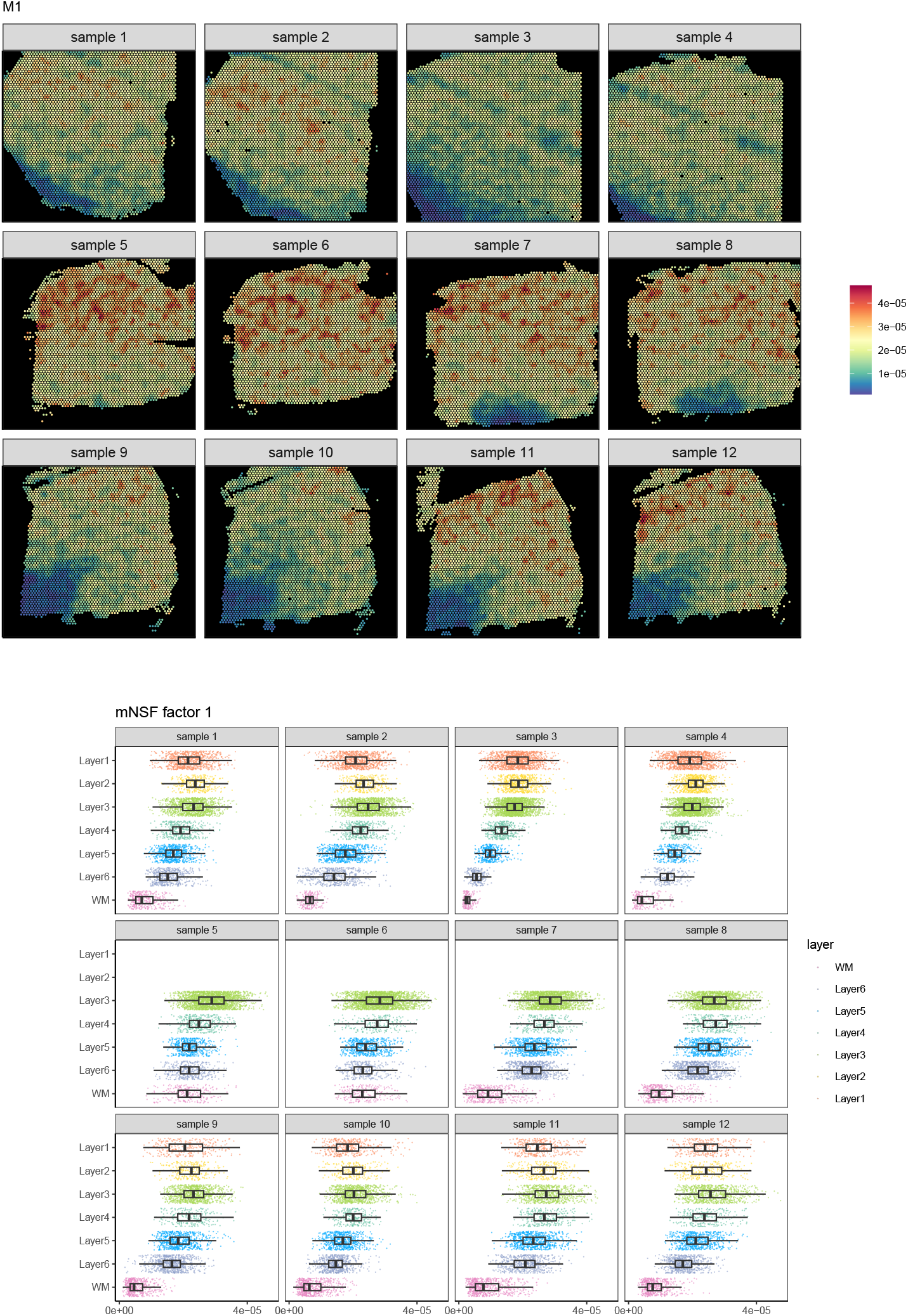
The value of each mNSF factor M1 for each of the 12 samples in DLPFC data.

**Supplementary Figure S7.**
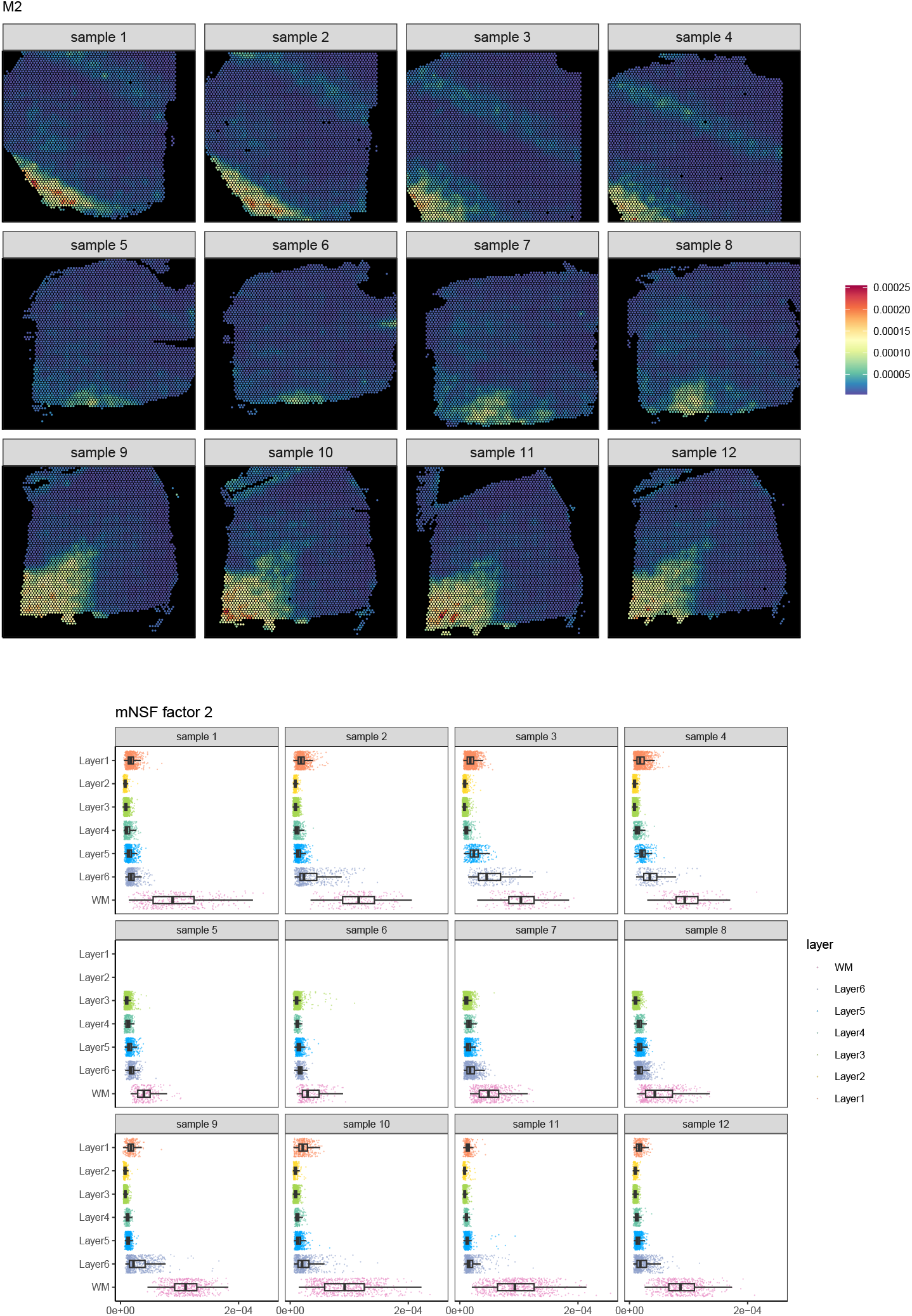
The value of each mNSF factor M2 for each of the 12 samples in DLPFC data.

**Supplementary Figure S8.**
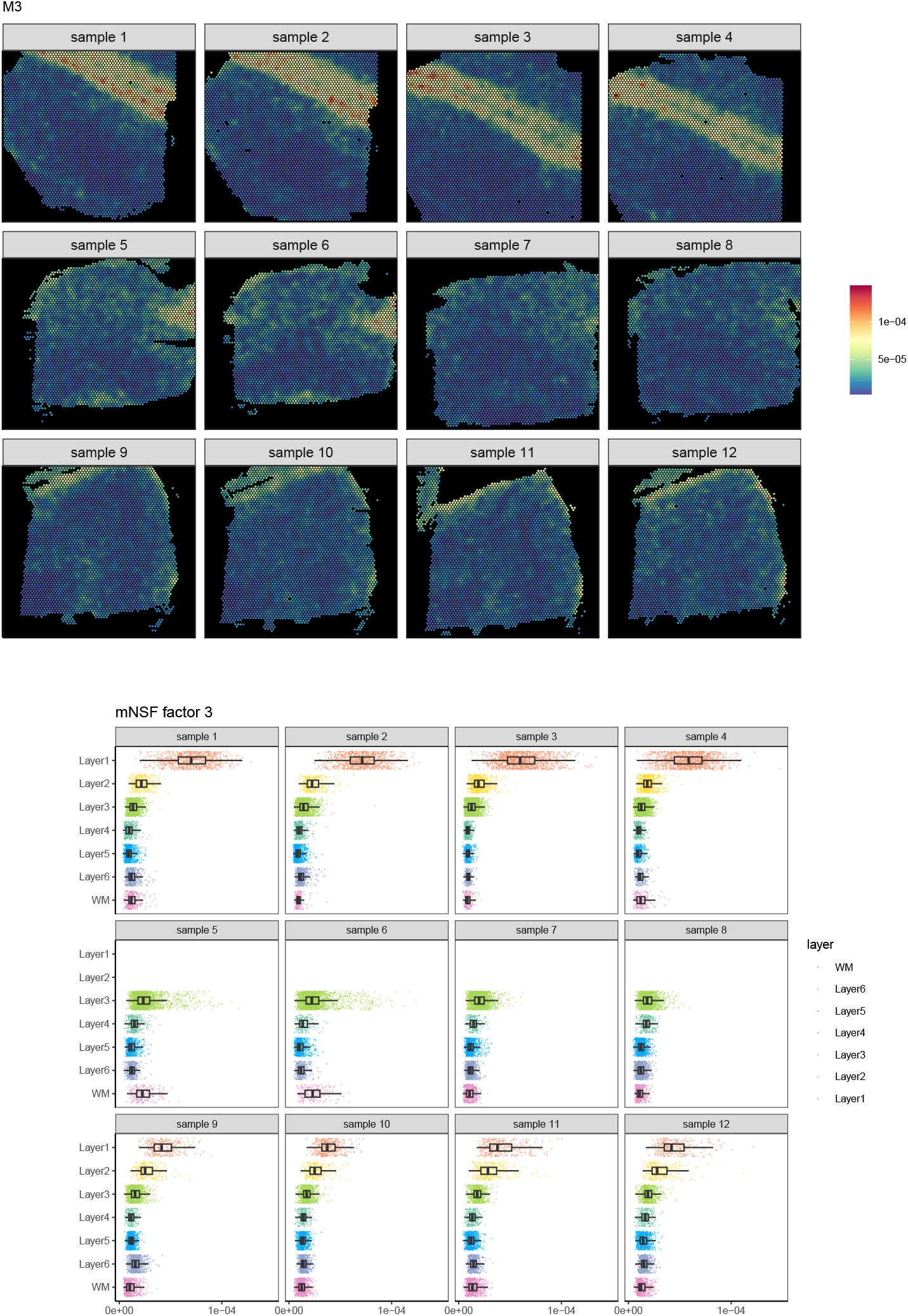
The value of each mNSF factor M3 for each of the 12 samples in DLPFC data.

**Supplementary Figure S9.**
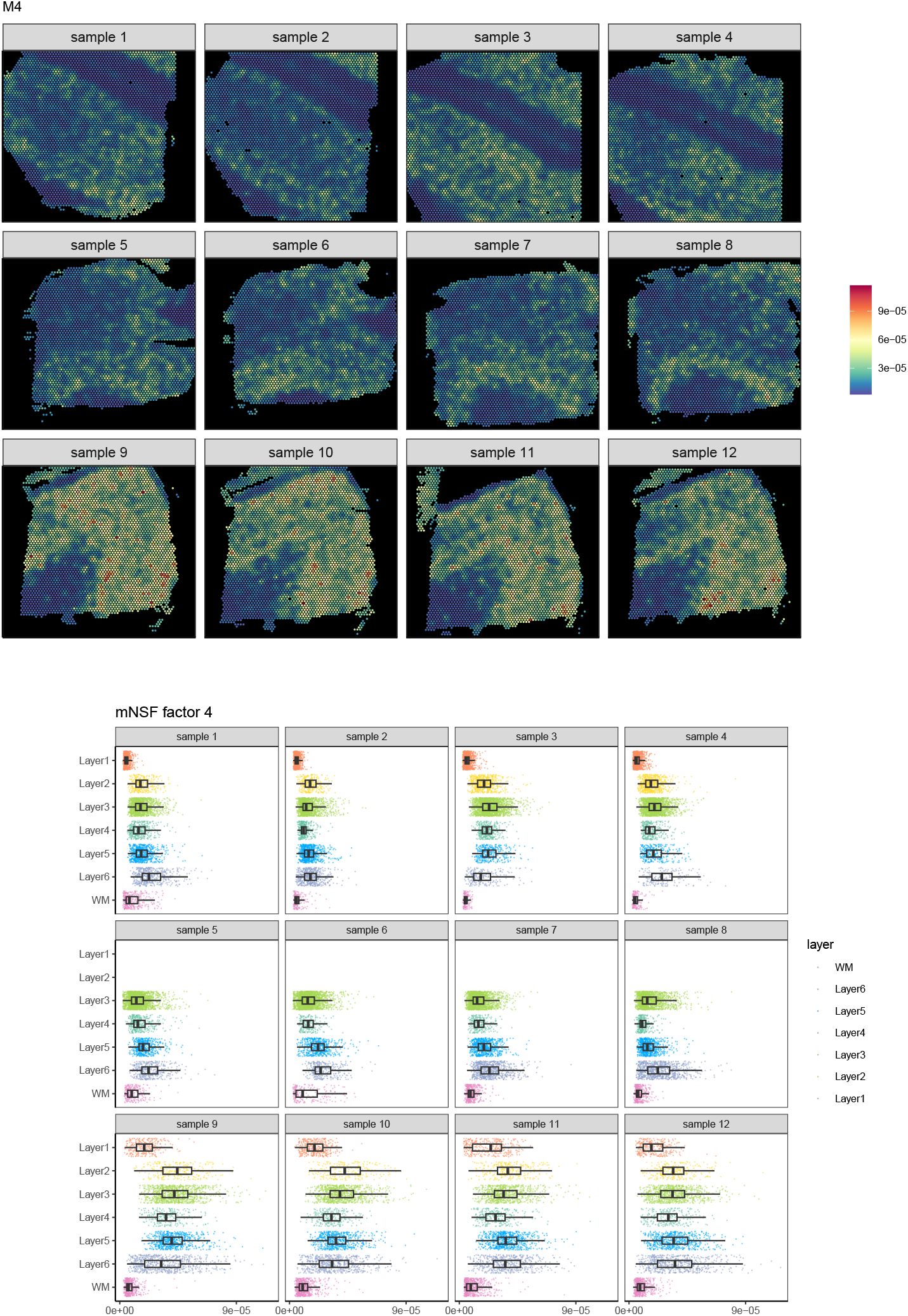
The value of each mNSF factor M4 for each of the 12 samples in DLPFC data.

**Supplementary Figure S10.**
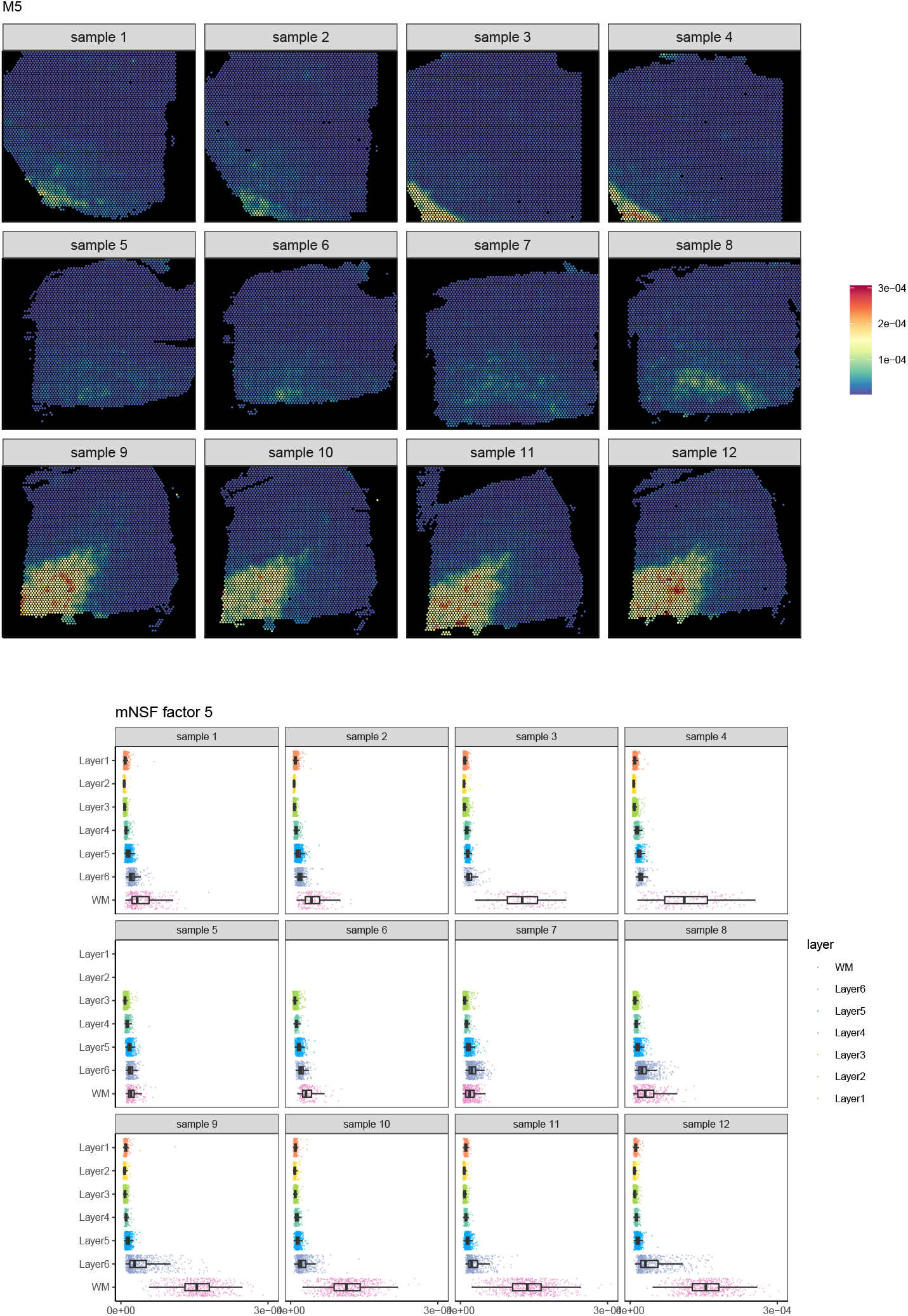
The value of each mNSF factor M5 for each of the 12 samples in DLPFC data.

**Supplementary Figure S11.**
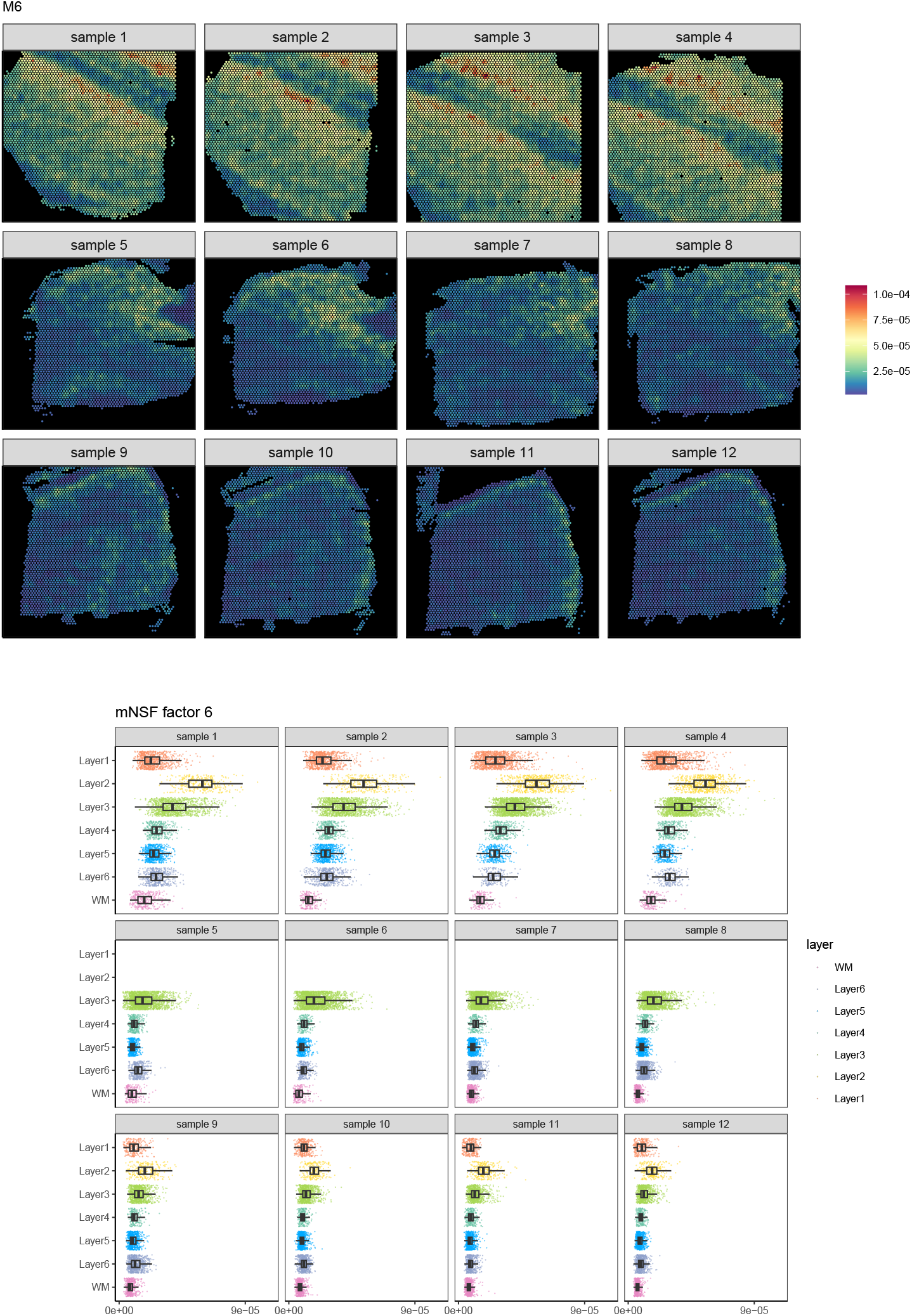
The value of each mNSF factor M6 for each of the 12 samples in DLPFC data.

**Supplementary Figure S12.**
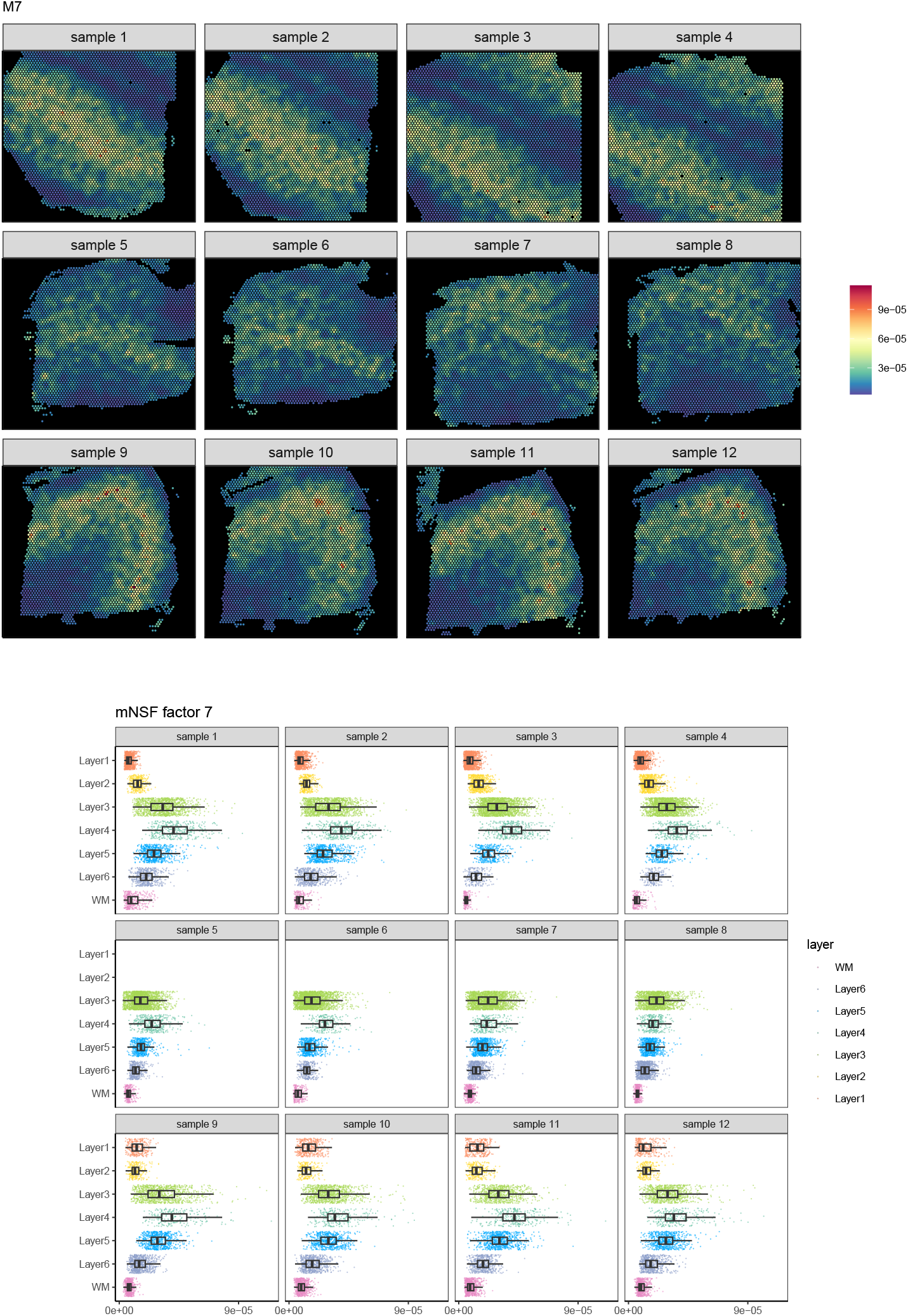
The value of each mNSF factor M7 for each of the 12 samples in DLPFC data.

**Supplementary Figure S13.**
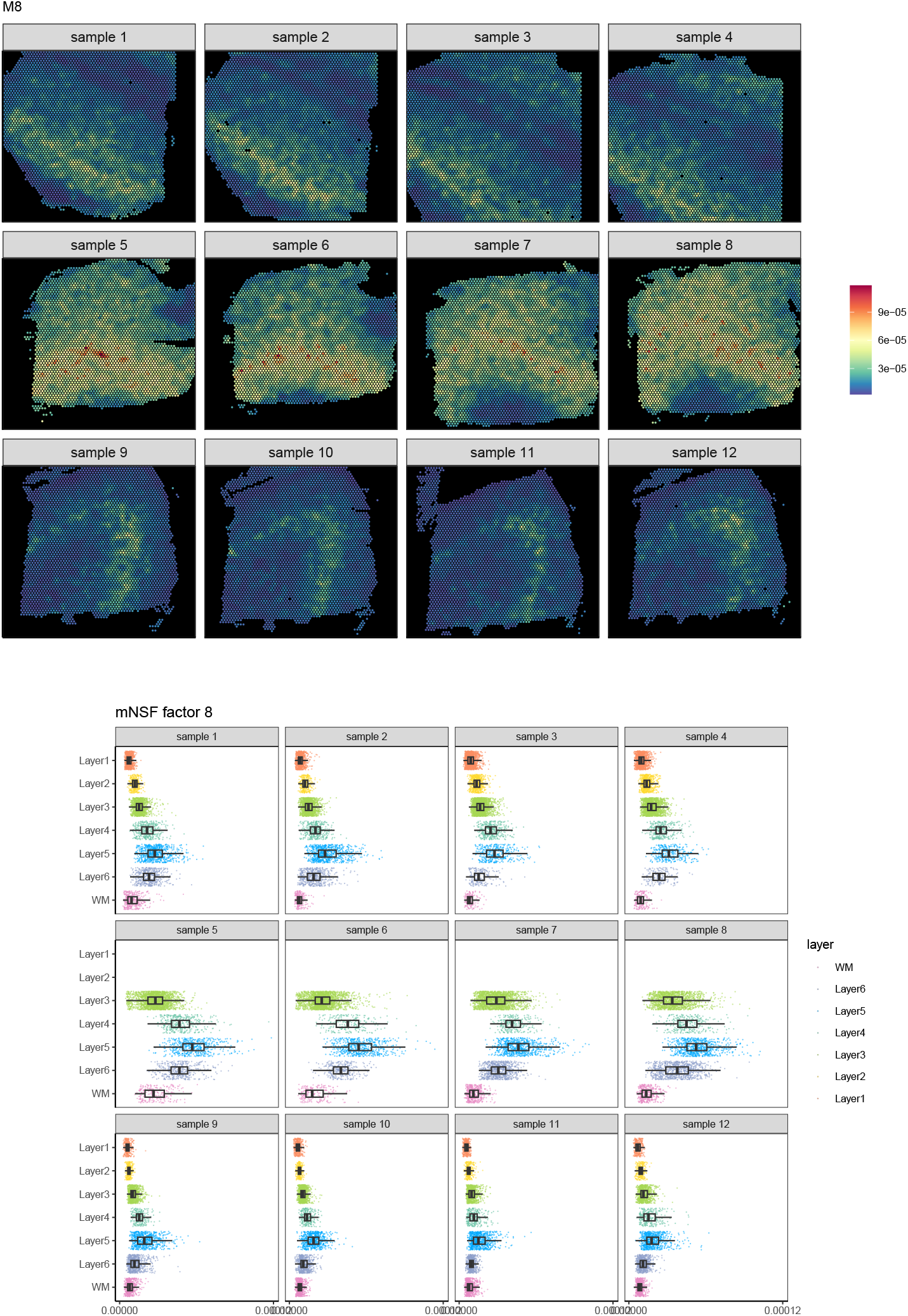
The value of each mNSF factor M8 for each of the 12 samples in DLPFC data.

**Supplementary Figure S14.**
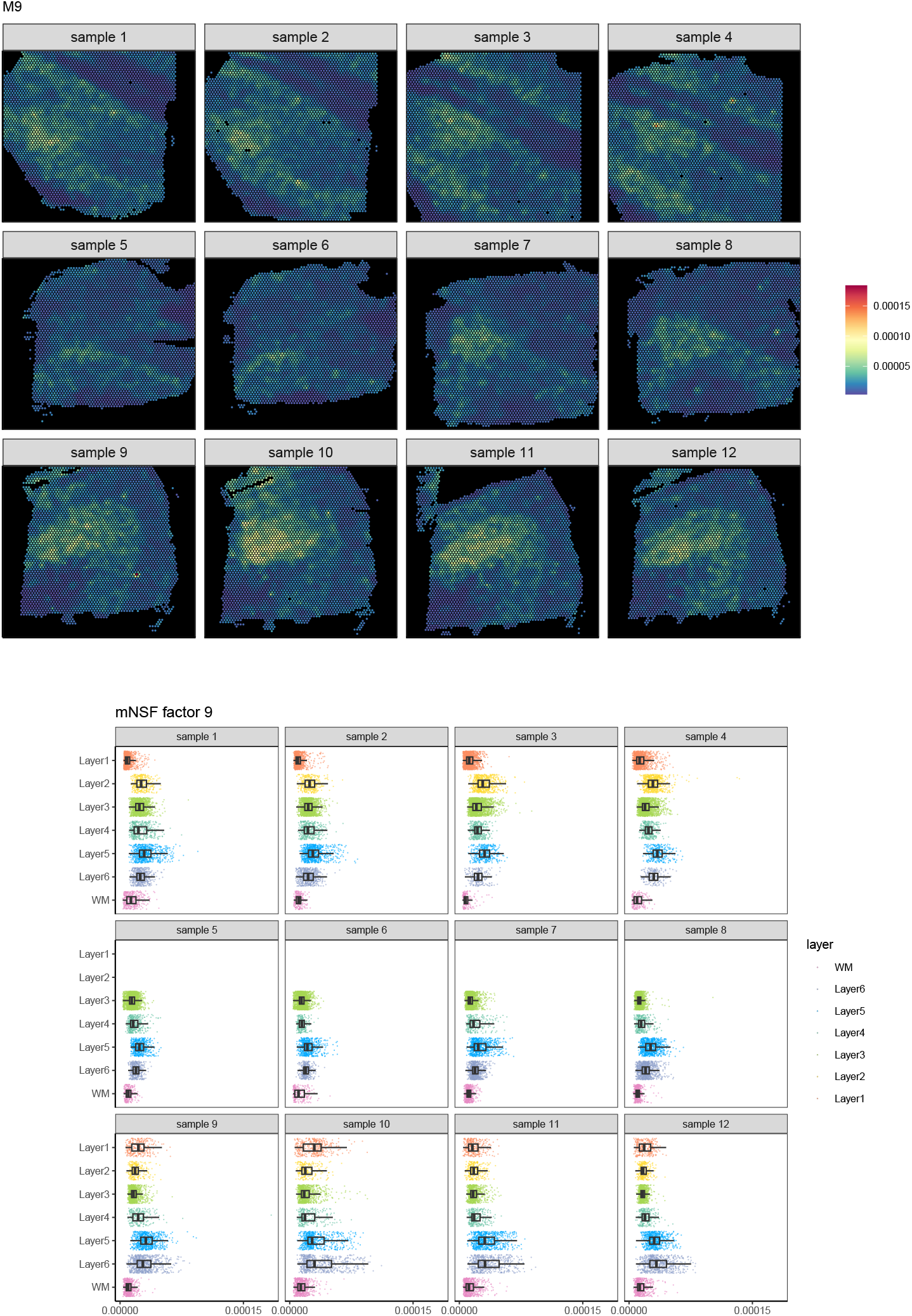
The value of each mNSF factor M9 for each of the 12 samples in DLPFC data.

**Supplementary Figure S15.**
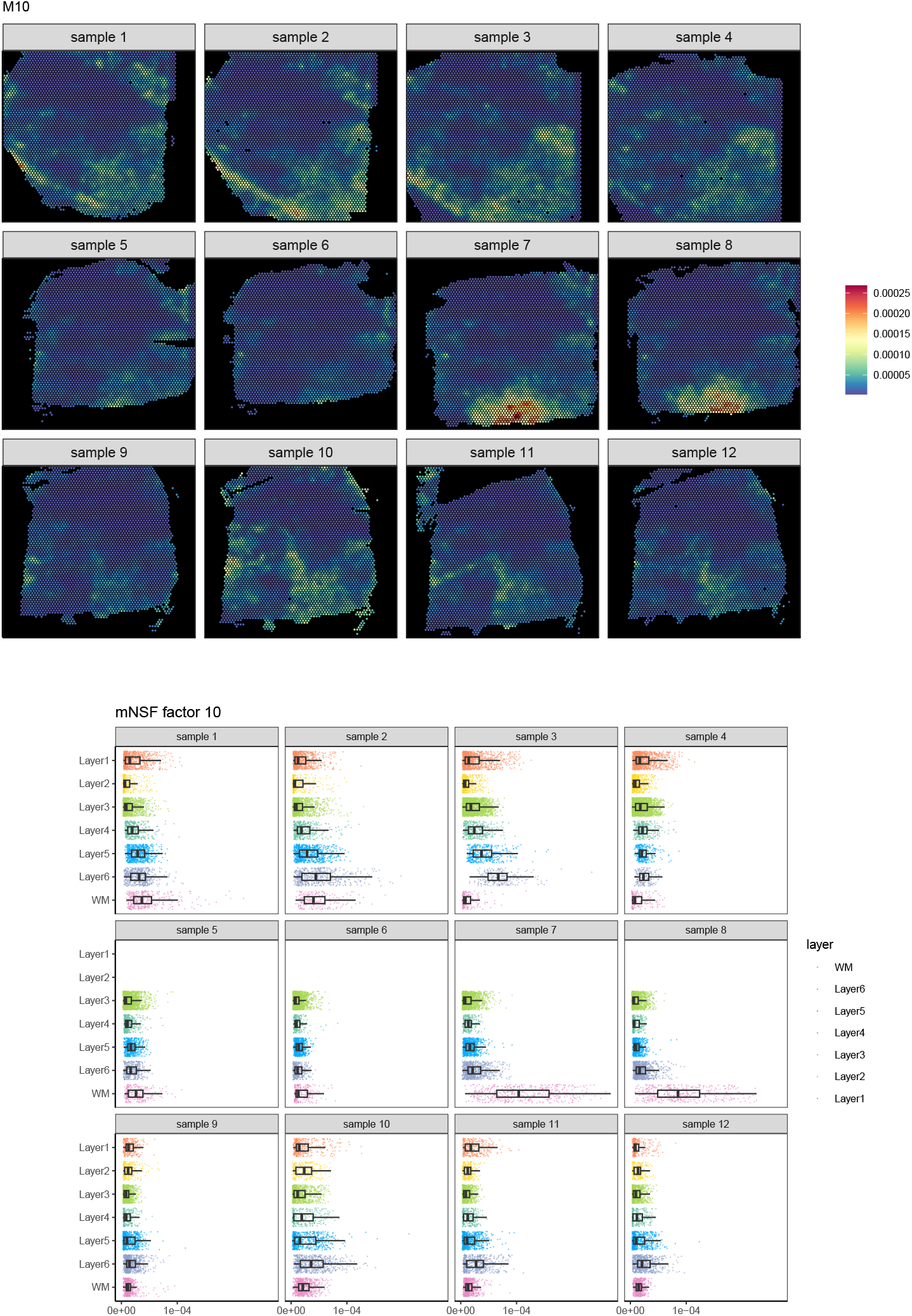
The value of each mNSF factor M10 for each of the 12 samples in DLPFC data.

**Supplementary Figure S16.**
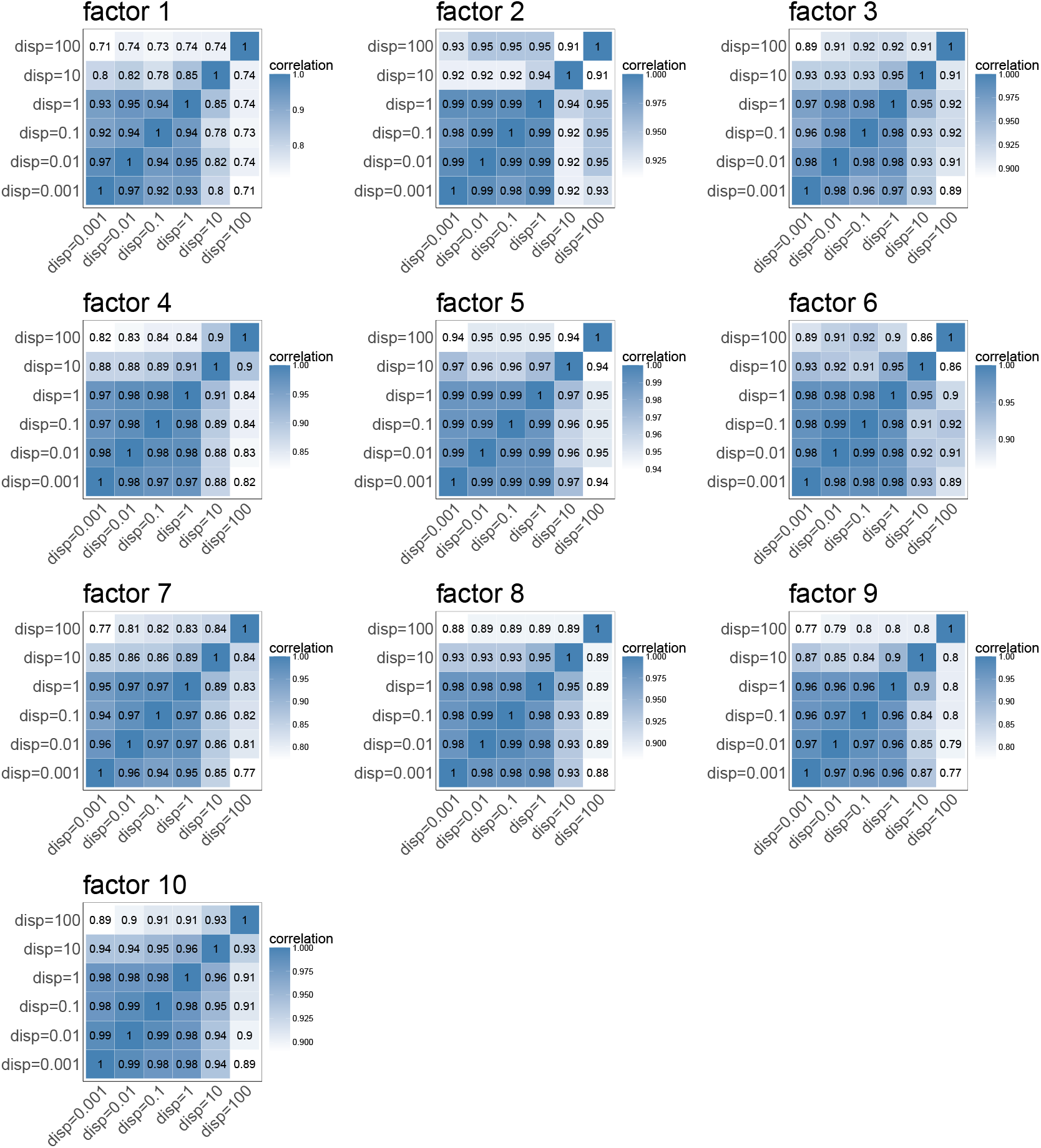
Sensitivity analysis of dispersion parameter on human DLPFC data. Correlation matrices showing the similarity between spatial factors obtained using different dispersion parameter values (ranging from 0.001 to 100) for each of the 10 factors identified in the DLPFC dataset. Each heatmap represents one factor, with colors indicating the correlation strength between factors obtained with different dispersion values. The high correlation values (> 0.8) observed across most parameter values for factors 1-10 indicate that mNSF results are generally robust to changes in the dispersion parameter within the range of 0.001 to 10, with some factors showing decreased correlation at dispersion=100.

**Supplementary Figure S17.**
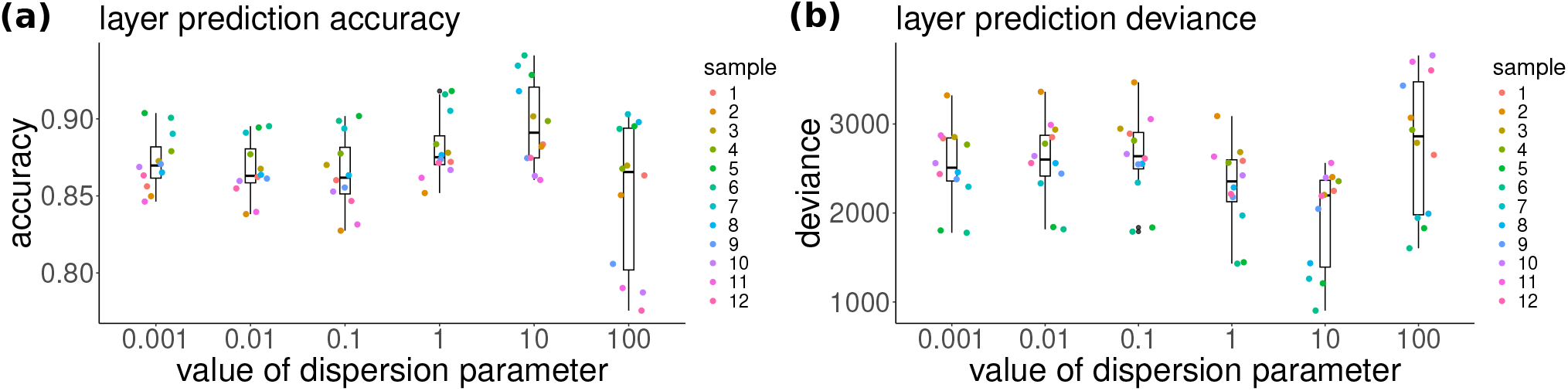
Performance of mNSF across dispersion parameter values on human DLPFC data. Analysis of how the dispersion parameter affects the association between mNSF factors and manually annotated cortical layers across 12 samples. We used a generalized linear model to predict layer and comapred to manual annotation on the training data. **(a)** Accuracy of our prediction. **(b)** Deviance measure for our prediction model (lower is better).

**Supplementary Figure S18.**
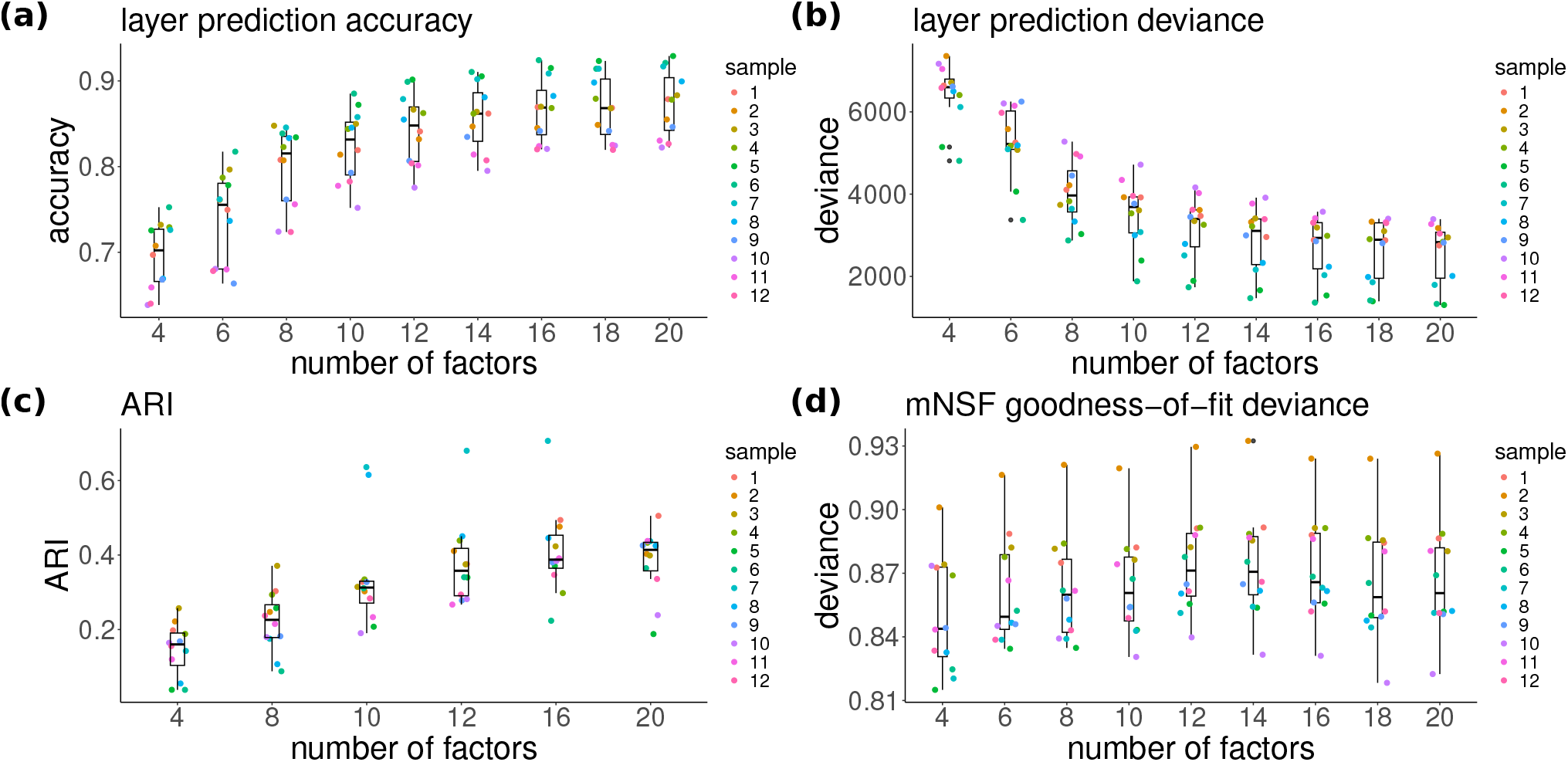
Evaluation of mNSF performance across different numbers of factors for DLPFC data. Systematic evaluation of mNSF performance using 4 to 20 factors across 12 samples. We used a generalized linear model to predict layer and comapred to manual annotation on the training data. **(a)** Accuracy of our model. **(b)** Deviance of our prediction model (lower is better). **(c)** Adjusted Rand Index (ARI): Quantifies the agreement between predicted spatial domains and known cortical layers. **(d)** mNSF goodness-of-fit deviance: measures the overall model fit using Poisson deviance between observed counts and predicted mean values.

**Supplementary Figure S19.**
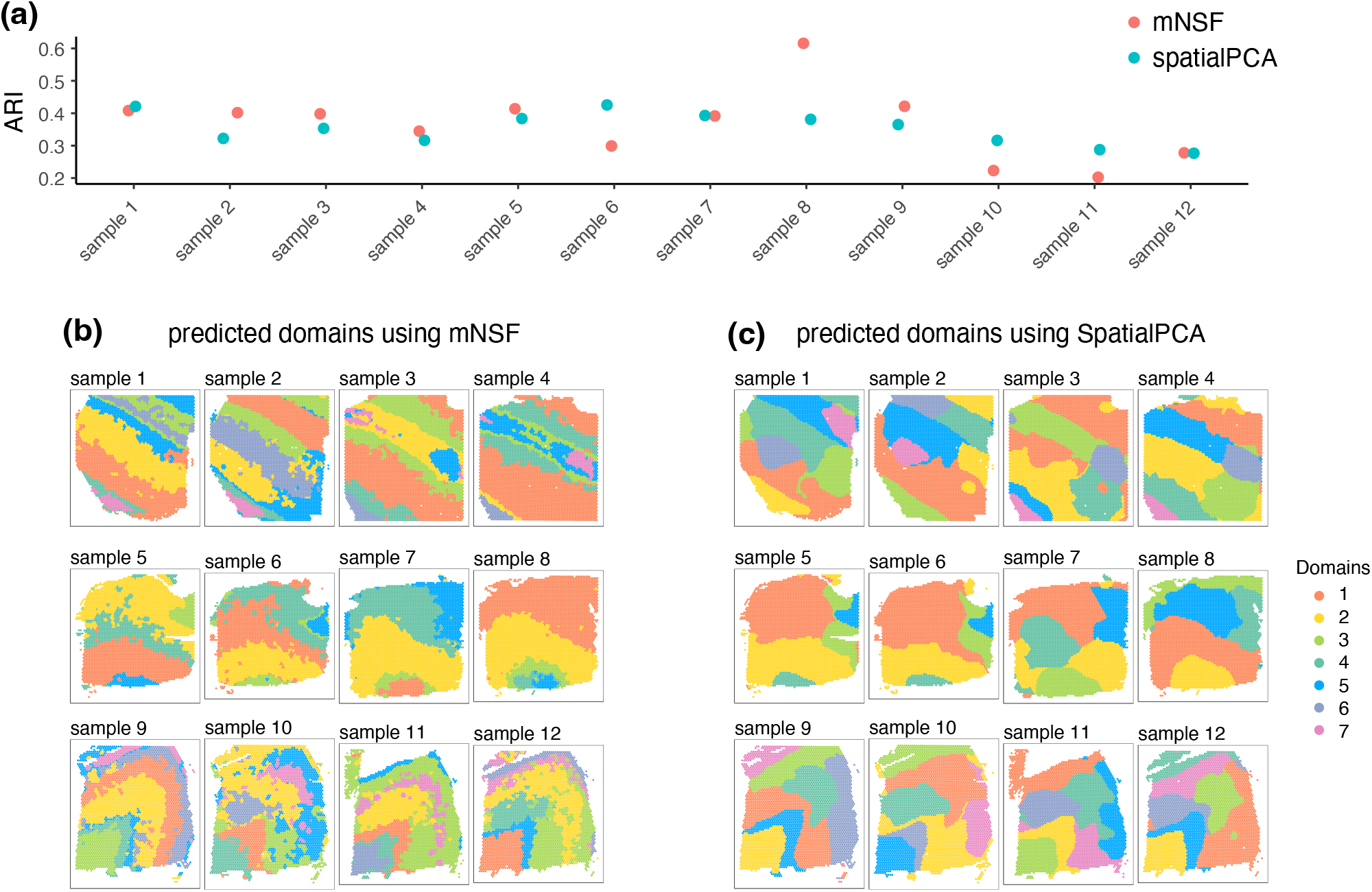
Comparison of domain identification performance between mNSF and SpatialPCA using the DLPFC dataset. **(a)** Adjusted Rand Index (ARI) scores comparing predicted domains to manually annotated layers across twelve samples. ARI values mostly ranged between 0.30 and 0.45, with neither method consistently outperforming the other across all samples. **(b)** Results of domain identification using mNSF, showing the predicted spatial domains for different samples. The colors represent distinct predicted domains within each sample. **(c)** Results of domain identification using SpatialPCA, showing comparable domain predictions for the same samples. The color scheme used is consistent within each method to facilitate comparison of domain structures.

**Supplementary Figure S20.**
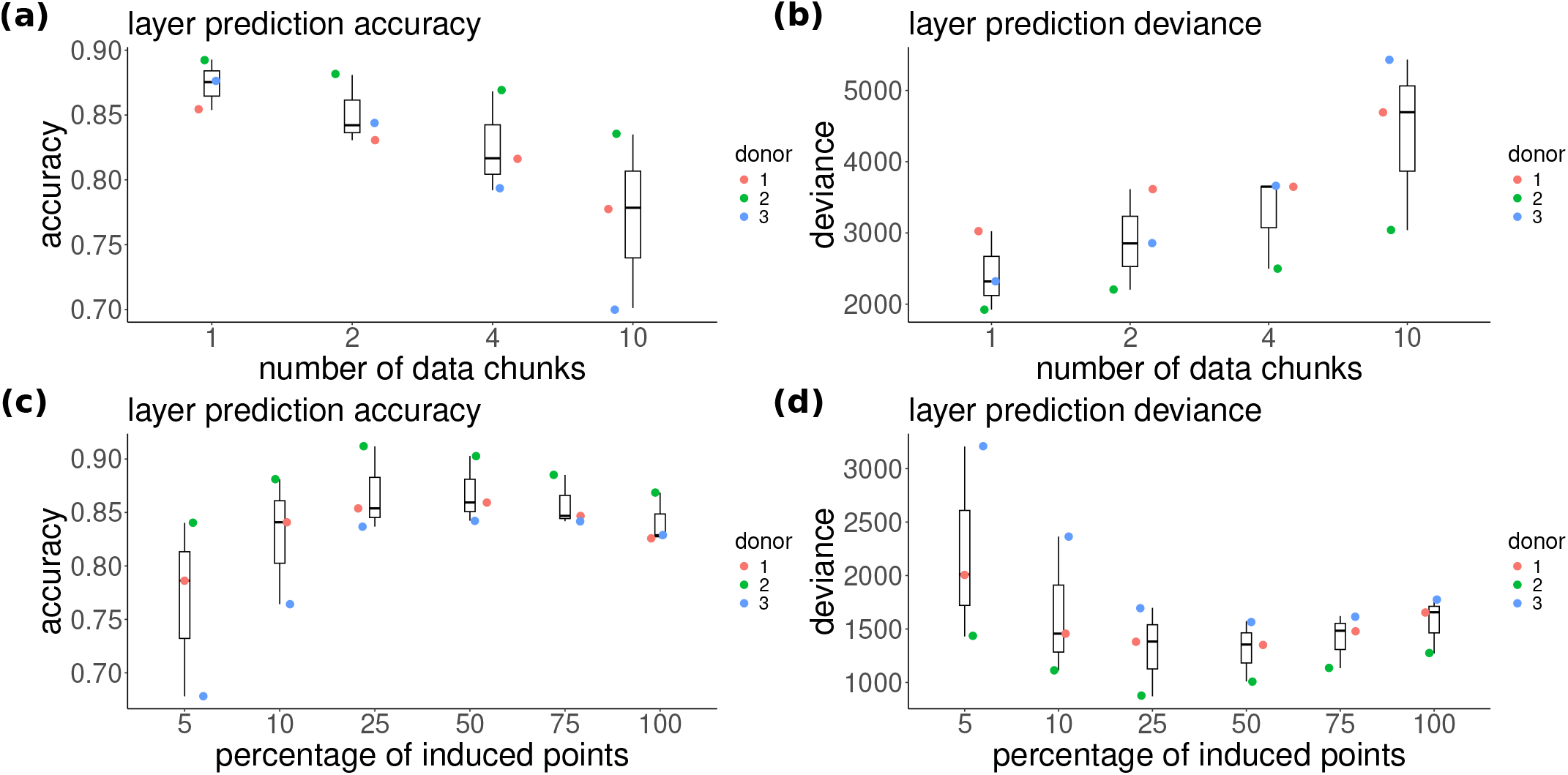
Performance evaluation of mNSF with varying induced points and data chunks on DLPFC data. Analysis of how induced points percentage and data chunking affect the association between mNSF factors and manually annotated cortical layers across three donors. **(a-b)** Layer prediction accuracy and deviance vs percentage of induced points. Higher accuracy and lower deviance indicate better performance. **(c-d)** Layer prediction accuracy and deviance vs number of data chunks.

**Supplementary Figure S21.**
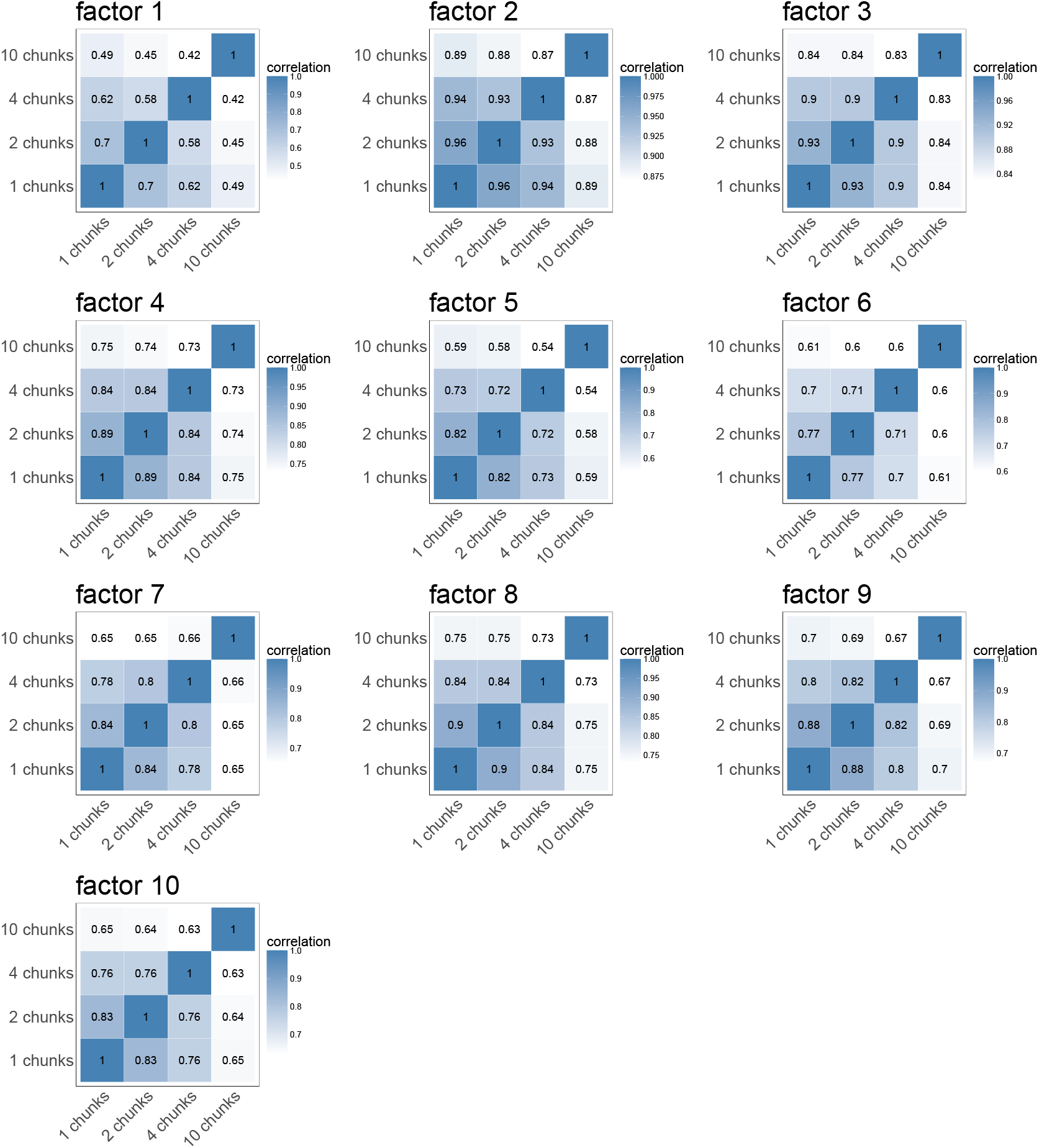
Impact of data chunking on factor correlation in DLPFC data. Correlation matrices showing the similarity between spatial factors obtained using different numbers of data chunks (1, 2, 4, and 10 chunks) for the 10 factors identified in the DLPFC dataset. Each heatmap represents one factor, with colors indicating the correlation strength between factors obtained with different chunk numbers. Lower correlations at higher chunk numbers suggest some loss of factor consistency with increased chunking.

**Supplementary Figure S22.**
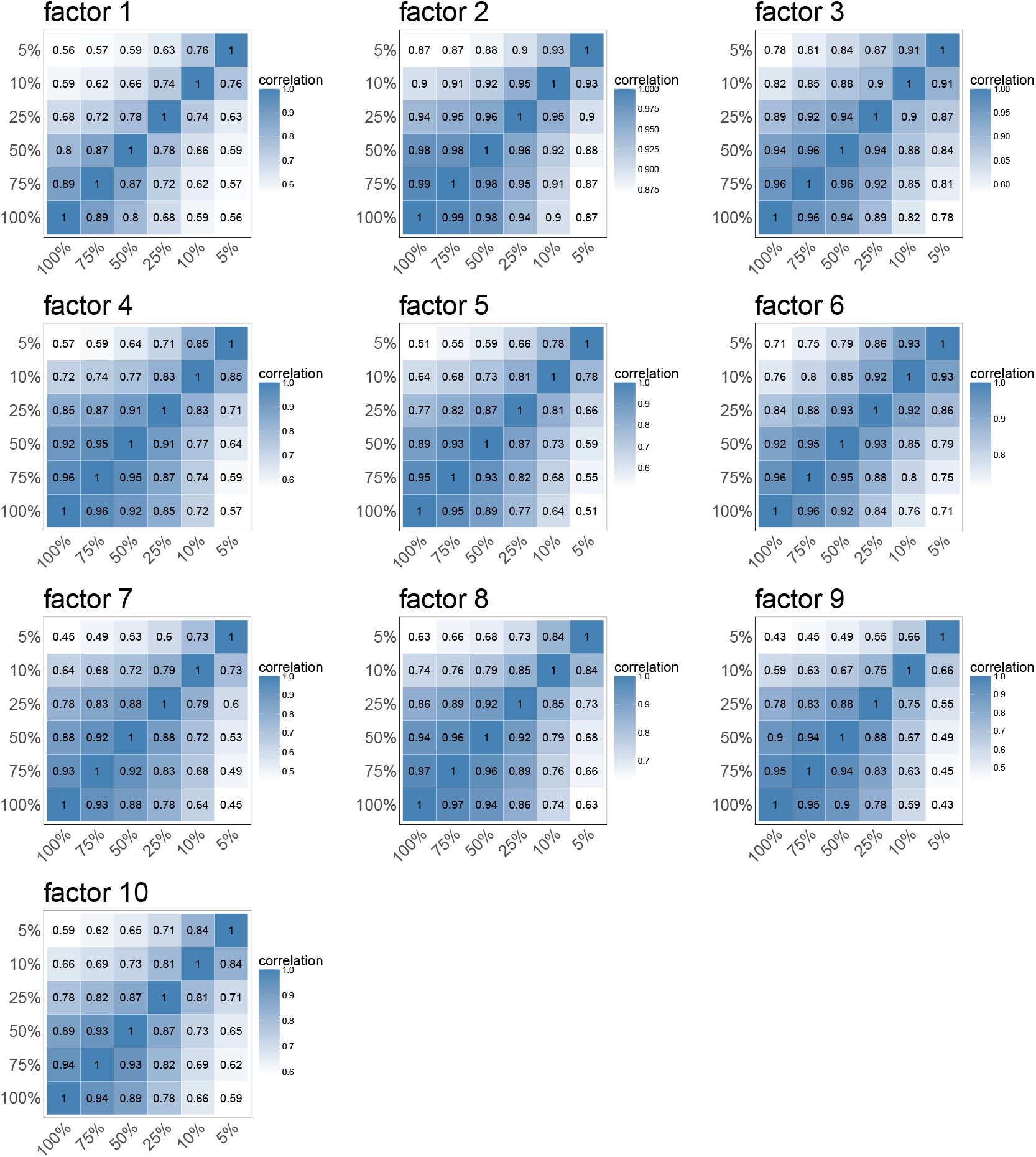
Effect of induced points percentage on factor stability in DLPFC data. Correlation matrices displaying the similarity between spatial factors obtained using different percentages of induced points (ranging from 5% to 100%) for factors 1-10. Each heatmap represents one factor, with correlation values indicated by color intensity. The high correlation values (generally > 0.8) between most percentage levels demonstrate that mNSF produces stable results across a broad range of induced point percentages, though some factors show decreased correlation at lower percentages.

## Notes

### Competing Interest Statement

The authors have declared no competing interest.

### Summary of Updates

We added work on - memory and runtime profiling - a cross technology analysis - comparison to SpatialPCA - More simulations

